# Deep Learning of Functional Perturbations from Condensate Morphology

**DOI:** 10.1101/2025.08.18.670955

**Authors:** Anita Donlic, Troy J. Comi, Sofia A. Quinodoz, Nima Jaberi-Lashkari, Krist Antunes Fernandes, Lifei Jiang, Lennard W. Wiesner, Ai Ing Lim, Clifford P. Brangwynne

## Abstract

Biomolecular condensates compartmentalize the interior of living cells to spatiotemporally organize complex functions, yet linking molecular interactions within condensates to their mesoscale organization remains a major challenge. To bridge this gap, we developed a neural network-based framework - Deep-Phase - that uses microscopy images to quantitatively classify condensate morphology changes resulting from pharmacological alterations in associated biochemical processes. We use Deep-Phase to precisely quantify time- and concentration-dependent structural perturbations to the multiphase nucleolus and show that they are tightly coupled to potencies of drugs inhibiting rRNA transcription and processing. Applying Deep-Phase in a chemical screen, we identify a unique nucleolar morphology and discover a role for a DNA topoisomerase in rRNA processing. Mechanistic studies of this morphology provide insights into how interfaces between nucleolar subcompartments are maintained. We demonstrate Deep-Phase’s adaptability to diverse cell lines, labels, and condensates, offering a powerful platform for uncovering cellular organizing principles and therapeutic targets.

**Highlights:**

- A deep learning framework, Deep-Phase, classifies and quantifies drug-induced changes in morphologies of nucleoli, nucleolar speckles, and viral cytoplasmic condensates, directly from images.
- Time- and concentration-dependent morphological responses to perturbation predict associated disruptions in RNA transcription and processing.
- Using Deep-Phase in a high-content small molecule screen reveals a unique nucleolar morphology induced by TOP1 inhibition.
- TOP1 inhibition leads to reduced levels and processing of large ribosomal subunit precursors and provides a mechanism for maintenance of nucleolar phase boundaries.

## Introduction

Spatial compartmentalization of biomolecular interactions is key to achieving complex cellular functions. Membraneless biomolecular condensates represent an important class of compartments that play diverse roles to control processes ranging from gene expression to RNA homeostasis^1–5^. Focused efforts spanning theoretical, computational, and experimental studies have identified molecular features of proteins and nucleic acids that drive the formation of condensates with richly textured multi-component and multi-phase organization^6–8^. However, it remains challenging to directly connect this mesoscale structural organization to the underlying biochemical activities within condensates, limiting our ability to extract mechanistic insights. Robustly and quantitatively linking nanoscale biochemical changes to mesoscale condensate structure is essential for enabling both the discovery of new emergent organizing principles and the identification of novel biomarkers and targetable mechanisms for disease treatment.

Quantitative imaging of biomolecular condensates has been foundational for the discovery of important principles of intracellular phase transitions and their functional relevance^9–14^. For example, imaging has been instrumental in mapping phase diagrams of biomolecular condensates, both in reconstituted *in vitro* systems with purified components^15–17^ and in cells^18–21^ as well as for uncovering principles of molecular partitioning into these bodies^10,22,23^. Compared to bulk biochemical readouts, imaging assays additionally enable single-cell resolution of phenotypes of interest while preserving spatial and morphological context. These capabilities enable a more nuanced assessment of cellular responses to genetic or pharmacological perturbations, which is important for developing a mechanistic understanding of cellular physiology and disease.

Condensate imaging-based approaches are particularly appealing in the context of pathologies, as disruption of condensate organization is increasingly recognized across a wide spectrum of diseases - including cancer, viral infections, and neurodegeneration^24–29^. This growing recognition has elevated condensates as attractive therapeutic targets, prompting efforts to identify small molecules that can selectively modulate their composition, dynamics, and function^30–33^. However, the field still lacks robust tools for systematically discovering such modulators. High-content, imaging-based screens (HCS) have emerged as a promising strategy^34,35^, but existing methods largely rely on pre-defined, low-dimensional image features (e.g. number, size, shape, partitioning) that provide limited insight into the underlying biochemical activities driving mesoscale condensate behavior. The use of these *a priori* defined metrics of interest carries the risk of overlooking other relevant impacts or even novel changes that could provide important biological insights^36^. Finally, such approaches lack the sensitivity needed to use images for direct measurements of biochemical potency, as is common in simplified biochemical or molecular binding assays.

Advances in deep learning algorithms for images, often called “computer vision”, are proving helpful by providing a quantitative but unbiased analysis approach for studying subcellular organization^37^, including condensates^38,39^. Deep learning methods have reinvigorated interest in HCS of both genetic and chemical libraries, leading to discoveries of fundamental biology^40,41^ and of therapeutic targets^42^. These algorithms employ multi-layered neural network architectures that progressively extract hierarchical image features, allowing the model to learn complex patterns of cellular organization directly from pixel data, without requiring predefined metrics of interest^43,44^. Convolutional layers enable detection of spatial relationships and fine-grained morphological structures, which could be especially helpful for studying condensates, given the diversity of their morphological and functional manifestations.

The eukaryotic cell contains a number of condensates that are particularly attractive for such deep-learning-based approaches, due to their large sizes and direct connections to fundamental processes guiding RNA homeostasis. Among nuclear condensates, the nucleolus is the largest, and exhibits a distinct multiphase organization that facilitates the complex and essential process of ribosome biogenesis, within its concentric phases (Figure 1A)^10,45,46^. Specifically, the innermost fibrillar center (FC) contains ribosomal RNA (rRNA) transcriptional machinery, the dense fibrillar component (DFC) is enriched in enzymes necessary for rRNA processing and small ribosomal subunit assembly, and the outermost granular component (GC) houses proteins that facilitate large ribosomal subunit and ribosomal particle formation. Importantly, previous studies have shown that inhibition of rRNA transcription and processing coincides with distinct changes in nucleolar integrity, suggesting that the maintenance of nucleolar structure depends on its function in ribosome biogenesis^47–50^. Changes in nucleolar morphology have thus generated interest for their use as HCS readouts to identify both genetic and chemical regulators of ribosome biogenesis^51–56^. While these studies established the nucleolus as a reliable indicator of cellular dysfunction, their reliance on traditional image analysis approaches measuring low-dimensional, predetermined features neglect the structural complexity of the nucleolus, and the intricate process of ribosome biogenesis within it. There is thus still major untapped potential for using unbiased deep learning-based approaches to quantitatively reveal condensate structure-function relationships, particularly for the nucleolus.

**Figure 1.**
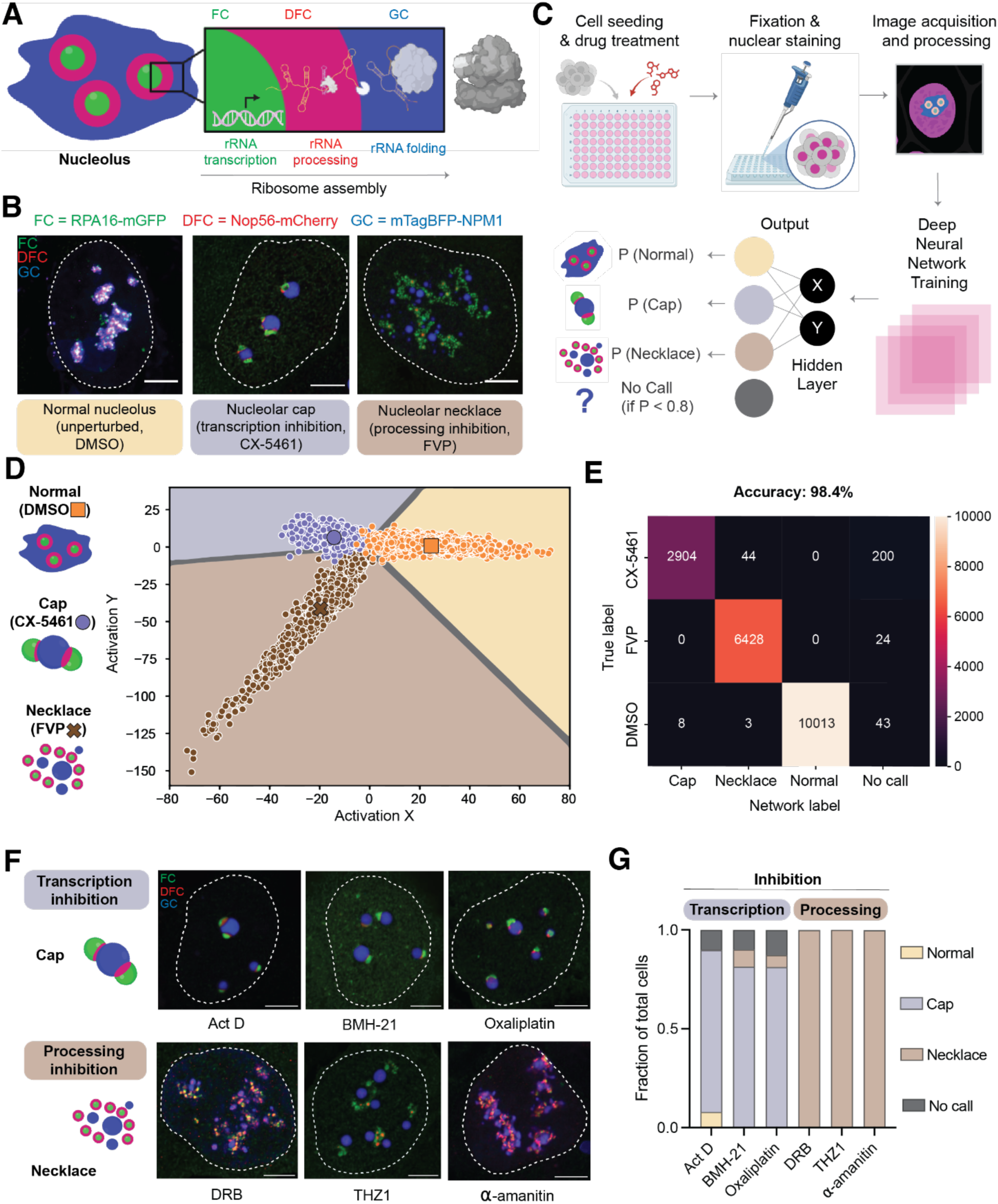
Deep-Phase accurately classifies distinct perturbations of nucleolar processes and morphology from fluorescent microscopy images. (A) Schematic of mammalian cell nucleolus displaying three compartments that organize ribosome biogenesis. The fibrillar center (FC) contains ribosomal RNA (rRNA) transcriptional machinery, the dense fibrillar component (DFC) harbors proteins necessary for rRNA processing and small subunit assembly, and the granular component (GC) houses ribosomal and non-ribosomal proteins that facilitate large subunit assembly and pre-ribosome formation. (B) Representative confocal microscopy images display U-2 OS cells expressing three nucleolar markers: FC, green, RPA16-mGFP; DFC, red, Nop56-mCherry; and GC, blue, mTagBFP-NPM1. Compared to the normal nucleolus (left, 0.25% DMSO), inhibition of rRNA transcription with 10 µM CX-5461 (middle) or 6.25 µM flavopiridol (FVP, right) for 2 hours causes the formation of canonical nucleolar “cap” or “necklace” phenotypes, respectively. Scale bar = 5µm. (C) Deep-Phase workflow of “training set” drug treatments (DMSO, CX-5461, FVP) in the three-color nucleolar osteosarcoma (U-2 OS) cell line with an automated image acquisition and analysis pipeline. Following drug treatment as described in (B), fixation and nuclear staining, cells are subjected to automated imaging, processing and augmentation before being presented to the neural network. The network classifies individual cells as having normal, cap, or necklace nucleolar morphology after calculating a pseudo-probability using Softmax function. If a cell’s probability of being in one of the three classes is below 0.8, it is classified as a “No Call”. (D) The hidden layer from the neural network provides X and Y coordinates for graphing network outputs in a two-dimensional “activation space”. Averages of cells (black borders) as well as individual cells (white borders) are placed in their expected regions. (E) Confusion matrix of cells from the test set displays high accuracy for classifying treatments into their corresponding nucleolar morphology class. (F) Representative confocal images and schematics of cap and necklace phenotypes caused by additional drug perturbations. All cells were treated for 2 hours, except for alpha-amanitin (4 hours). Act D = Actinomycin D (0.001 µM), BMH-21 (1µM), Oxaliplatin (25 µM), DRB = 5,6-dichlorobenzimidazole riboside (25 µM), THZ1 = (0.5 µM), Alpha-amanitin (6.25 µg/mL). Scale bar = 5µm. (G) Fractions of cells from treatments in (F) as classified by the neural network and calculated by the Softmax function. *n* ≥ 500 cells per condition.

Here, we develop a deep neural network-based approach, Deep-Phase (Deep learning of Phase-separated condensates) that can accurately classify and quantify specific condensate disruptions, and use it to discover a new regulator of nucleolar form and function. Deep-Phase extracts complex condensate features directly from fluorescence microscopy images of human cells, and is generalizable across diverse condensate types. We quantify distinct changes in the morphology of nucleoli, nuclear speckles, and viral cytoplasmic condensates, as a function of drug treatment time and concentration. The extent of morphology alterations can be used to generate dose-response curves that are consistent with literature IC_50_ values from biochemical experiments, and predictive of the degree to which RNA-dependent processes are disrupted. We utilize Deep-Phase in an RNA-targeted small molecule screen, identifying a previously unrecognized role for the DNA topoisomerase TOP1 in rRNA processing, which Deep-Phase detects through a distinct nucleolar morphological fingerprint. We show that TOP1 inhibition, unlike other perturbations, disrupts global nucleolar organization while preserving nucleolar phases, by limiting the buildup of misprocessed rRNA species. Together, these findings demonstrate that integrating quantitative deep learning with functional condensate studies serves as a powerful discovery tool for uncovering new insights into the interplay between condensate structure, function, and therapeutic targeting.

## RESULTS

### Deep-Phase accurately classifies nucleolar morphologies induced by distinct perturbations of ribosome biogenesis steps

We first sought to develop Deep-Phase for classifying the morphology of the multiphase nucleolus, as one particularly prominent condensate central to cellular physiology and disease. We chose to image nucleoli in human osteosarcoma cells (U-2 OS) because of their clinical relevance, since they are patient-derived, as well as amenability to automated microscopy of intracellular structures, due to their large size and flat morphology. We incorporated fluorescent protein markers of all three nucleolar compartments by stably expressing exogenous proteins that localize to these regions. Specifically, the FC was visualized with the RNA Polymerase I (Pol I) subunit protein RPA16 fused to monomeric green fluorescent protein (mGFP), the DFC was visualized with Nop56, a core component of the ribosomal small subunit (SSU) processome^57^ fused to mCherry, and the GC was labeled with monomeric blue fluorescent protein (mTagBFP2)-fused nucleophosmin (NPM1), an abundant GC protein that assists in the maturation and flux of ribosomal particles by directly interacting with large ribosomal subunit precursors^58^. Using antibody staining of wild-type U-2 OS cells, we verified that the nucleolar morphologies are similar to our tagged cell line, hereafter referred to as the three-color nucleolar line (Figure 1B left and Figure S1A).

We additionally confirmed that the three-color nucleolar line displays canonical phenotypes upon disruption of rRNA transcription and processing. Specifically, we used the selective Pol I inhibitor CX-5461^59,60^, and observed the formation of the well-known nucleolar “cap” morphology^61,62^ within 2 hours (Figure 1B, center). We also subjected cells to flavopiridol (FVP), an inhibitor of cyclin-dependent kinases (CDKs)^63^ that indirectly alters nucleolar function by inhibiting synthesis of Polymerase II (Pol II)-transcribed RNA involved in ribosome biogenesis^48,64–66^. Two-hour FVP treatment gave rise to the expected nucleolar “necklace” morphology (Figure 1B, right panel). We next utilized the three-color nucleolar line treated with either CX-5461 or FVP, as a training set to establish an automated microscopy-based Deep-Phase pipeline. Following cell treatment, imaging, and processing (Figure 1C, top panel, also see Methods), this approach yielded tens of thousands of cell images from each treatment category. These images were then used to train convolutional neural networks (CNNs) with different numbers of convolutional layers, all of which are built from an underlying ResNet architecture^67^, one of the most established and benchmarked architectures for image analysis^68,69^. By assessing the loss function, we found that the ResNet50 CNN had the highest performance for both the training set and the test set that the network had not previously seen (Figure S1B). To enable visualization of individual cell classification distributions within each treatment category, we incorporated a hidden encoder-decoder layer prior to the final fully connected layer. This layer collapses the extracted features into two values (X and Y), which are linearly connected to the three class outcomes (Figure 1C, bottom panel), enabling us to assign each cell with an X and Y coordinate and plot it in a 2-dimensional activation space (Figure 1D). We found that this trained network yielded higher than 98% accuracy; most of the predicted network calls for different conditions from the test set matched the true condition label as visualized by the confusion matrix (Figure 1E).

We next examined whether Deep-Phase could similarly detect phenotypes from other drugs that disrupt rRNA transcription and processing through RNA polymerase inhibition. We treated cells with Actinomycin D, BMH-21, and oxaliplatin that are known to inhibit rRNA transcription^48,49,70^ as well as DRB, THZ1, and alpha-amanitin that are known to inhibit rRNA processing via Pol II transcriptional modulation^48,64,71–73^. We verified that these transcription and processing inhibitors induced cap or necklace phenotypes, respectively (Figure 1F), and found that they were classified as expected by population proportions from Deep-Phase outputs (Figure 1G). Together, these results demonstrate that Deep-Phase can identify and classify nucleolar morphology perturbations with high accuracy and generalizability.

### Measured changes in nucleolar morphology by Deep-Phase are a direct and specific fingerprint of altered nucleolar functional pathways

The magnitude of a small molecule’s effect on its cellular target and associated functional outcome is time- and concentration-dependent, although extracting this information directly from cellular images is typically not possible. We therefore examined whether such effects could potentially be captured and quantified as gradual alterations in nucleolar morphology by Deep-Phase. Upon fixing cells in 30 minute intervals over 2 hours, a gradual change from an unperturbed interphase nucleolus to the cap morphology was indeed observed in cells treated with CX-5461 (Figure 2A, top); an equivalent transition to a necklace morphology was observed over 2 hours of FVP treatment (Figure 2A, bottom). Furthermore, we observed dose-dependent transitions in cap and necklace formation at the 2 hour time point when treating cells with increasing drug concentrations (Figure 2B). Specifically, with CX-5461 treatment, we observed that while the nucleolar GC layer appears to shrink and round at intermediate concentrations (0.125 – 1 µM), FC fusion and positioning on the GC surface (i.e. cap formation) only occurs at higher concentrations (5 and 10 µM) (Figure 2C, top and S2A, top). Similarly, treating cells with increasing concentrations of necklace-forming FVP showed that lower concentrations (0.05 – 0.1 µM) cause GC layer fragmentation, but GC rounding and further separation from FC and DFC layers that are characteristic of necklaces occur at higher concentrations (0.125 µM and higher) (Figure 2C, bottom and S2A, bottom). We quantified these changes by projecting a treatment vector defined by the change in average response at a given time or concentration onto a standard vector, drawn between the trained “normal” to the trained drug-perturbed average (Figure S2B). The magnitude of each projected vector is then defined as the “nucleolar morphology response”. Analyzing the results and plotting population average coordinates for each time frame in activation space revealed that CX-5461- and FVP-treated cells steadily moved from the “Normal” to “Cap” and “Necklace” space, respectively, either in time at a fixed concentration, or as a function of concentration at fixed time (Figure 2C-D). Analyzing the time-dependent magnitude of vector projections, we can measure morphology formation rates (Figure S2C), while analyzing the concentration-dependent magnitude of vector projections, we can plot dose-response curves and measure corresponding nucleolar morphology response concentrations (RC_50_). CX-5461 treatment provided an RC_50_ range from 0.28-0.38 µM (95% confidence interval, Figure 2E). FVP treatment resulted in an order of magnitude lower range (RC_50_ = 0.074-0.104 µM, Figure 2F) relative to CX-5461, which corresponds to nucleolar morphology changes being visible at lower FVP concentrations compared to CX-5461 (Figure 2B, 2D, and S2A, bottom). Additionally, we detected comparable nucleolar morphology changes in wild-type U-2 OS cells labelled with different nucleolar compartment markers via immunofluorescence, yielding values of RC_50_ comparable to those measured in the three-color overexpression line (Figure S1C-D). Together, these results illustrate how Deep-Phase enables integrated and unbiased extraction of trajectories through a complex morphological space as a function of drug concentration and treatment time.

**Figure 2.**
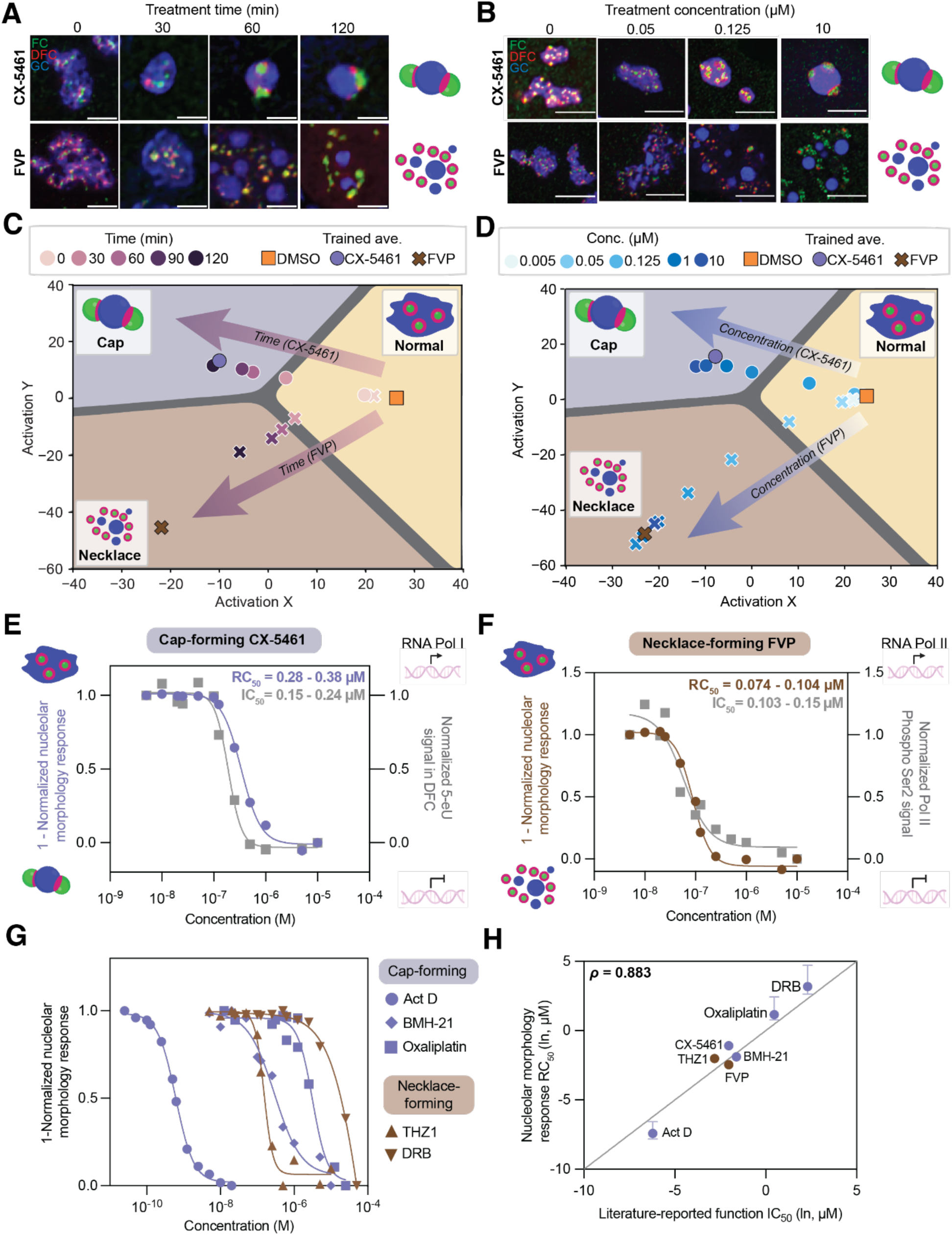
Deep Phase quantifies time- and concentration-dependent changes in nucleolar morphology, which reflect underlying functional disruptions. (A) Representative confocal microscopy images and schematics of nucleolar morphology change upon CX-5461 addition (10 µM, top) or FVP addition (6.25 µM, bottom) over different treatment times. Scale bar = 5µm. (B) Representative confocal microscopy images and schematics of nucleolar morphology change upon treatment with increasing concentrations of CX-5461 (top) or FVP (bottom) for 2 hours. Scale bar = 5µm. (C) Deep-Phase outputs of population averages for CX-5461-(10 µM) or FVP-treated (6.25 µM) cells in 30 minute intervals plotted on activation space. Trained class averages are used to generate vector projections for quantifying morphology changes (see Methods, Figure S2B). *n* ≥ 200 cells per time point per treatment condition. (D) Deep-Phase outputs of population averages for CX-5461- or FVP-treated cells at varying concentrations over 2 hours plotted on activation space. Trained class averages are used to generate vector projections for quantifying morphology changes (see Methods, Figure S2B). *n* ≥ 200 cells per concentration per treatment condition. (E) Dose-response curves for nucleolar morphology change generated from vector algebra done on treatment averages shown in (D) (left axis) and 5-eU intensity change (right axis, see Methods) for cap-forming CX-5461. Values were normalized to the lowest and highest treatment concentrations and curves were generated using a log-scaled variable slope least-squares fit. RC_50_ value ranges represent 95% confidence intervals. *n* ≥ 200 cells per concentration per condition. (F) Dose-response curves for nucleolar morphology change generated from vector algebra done on treatment averages shown in (D) (left axis) and phosphorylated Pol II antibody intensity change (right axis, see Methods) for necklace-forming FVP. Values were normalized to the lowest and highest treatment concentrations and curves were generated using a log-scaled variable slope least-squares fit. RC_50_ value ranges represent 95% confidence intervals. *n* ≥ 200 cells per concentration per condition. (G) Dose-response curves for a variety of other cap- and necklace-forming drugs (see Figure 1F-G). Values were normalized to the lowest and highest treatment concentrations and curves were generated using a log-scaled variable slope least-squares fit. RC_50_ values from curve fits are listed in Table S1-S2. *n* ≥ 200 cells per concentration per condition. (H) Tight correspondence between RC_50_ values from nucleolar morphology disruptions as measured by Deep-Phase and literature-reported IC_50_ values from functional readouts (see Table S1-S2). An average value is plotted for drugs with multiple literature-reported IC_50_ values. Error bars for RC_50_ values represent 95% confidence intervals. All measurements were ln-transformed before calculating the Spearman correlation coefficient (*ρ*) and a straight line was drawn through (-10,-10) for reference.

We next hypothesized that the morphological trajectories measured by Deep-Phase are reflective of underlying functional changes. To test this, we aimed to directly measure and compare the concentration-dependence of functional biochemical changes induced by these pharmacological perturbations. Specifically, to probe functional changes for cap-forming drugs such as CX-5461, we measured nucleolar rRNA transcriptional output by adapting a nucleotide analog incorporation paradigm, based on 5-ethynyluridine (5-eU) pulse-chase labeling^74–76^. Briefly, cells were fed with 5-eU, whose incorporation into nascent RNA transcripts can be visualized with a fluorophore using click chemistry, allowing us to quantify RNA transcriptional activity in the nucleolus (Figure S2D). The resulting functional dose-response curve obtained for CX-5461 was similar to the one obtained with the nucleolar morphology measurement by Deep-Phase; IC_50_ values ranged from 0.15-0.24 µM (95% confidence interval, Figure 2E) and were also in agreement with previously reported values using a different method (Table S1). For FVP-treated cells, we measured active RNA Pol II levels via immunofluorescence, as a proxy for Pol II phosphorylation by CDK9 (Figure S2E), which is inhibited by necklace-forming drugs^63,77^. Again, we obtained IC_50_ values consistent with previous reports using other approaches, and in agreement with the RC_50_ values measured by Deep-Phase (Figure 2F, Table S2). This striking ability of Deep-Phase to predict IC_50_ values through Deep-Phase RC_50_ measurements was also possible with other cap- and necklace-forming drugs (Figure 1F-1G), spanning a range of potency from low-nanomolar to high micromolar inhibitors (Figure 2G). All values were in agreement with previously reported functional IC_50_ measurements for these drugs (Figure 2H, Table S1-S2). As we observed a striking homogeneity of nucleolar morphology from different cells treated with the same drug concentration (Figure S2A), these results suggest that nucleolar organization is finely tuned to the levels of rRNA intermediates. We therefore conclude that the extent of distinct nucleolar morphology disruptions detected by Deep-Phase is a quantitative fingerprint of specific alterations in nucleolar biochemical activities, providing a powerful framework for linking mesoscale condensate organization to underlying molecular processes.

### Deep-Phase measures functional disruptions from mesoscale structure changes of nuclear speckles and cytoplasmic RSV inclusion bodies

We next sought to test if the Deep-Phase approach can be more broadly extended to other condensates that host important RNA regulatory processes. Nuclear speckles are hubs of messenger RNA splicing that form around transcriptionally active Pol II loci (Figure 3A). Previous studies reported that perturbing Pol II transcription or transcript splicing with small molecule inhibitors leads to speckle rounding or dilation, respectively^78–82^. We were able to reproduce these phenotypes in a HEK293 cell line containing an endogenously tagged nuclear speckle marker, SRRM2, using FVP and Pladienolide B (Plad B, Figure 3B). We then adapted the Deep-Phase neural network to analyze these images and create a “speckle activation space” (Figure 3C), confirming that these distinct changes can be detected with high accuracy (Figure S3A). Using a vector projection approach as described above (Figure S2B), we also measured dose-response curves of speckle morphology changes, upon treatment with FVP (RC_50_ = 0.083 - 0.092 µM, 95% confidence interval) and Plad B (RC_50_ = 0.0023 - 0.0048 µM, 95% confidence interval, Figure 3D and S3B). The quantified RC_50_ values matched function-related IC_50_ measurements reported previously (Figure 3D inset and Table S3).

**Figure 3.**
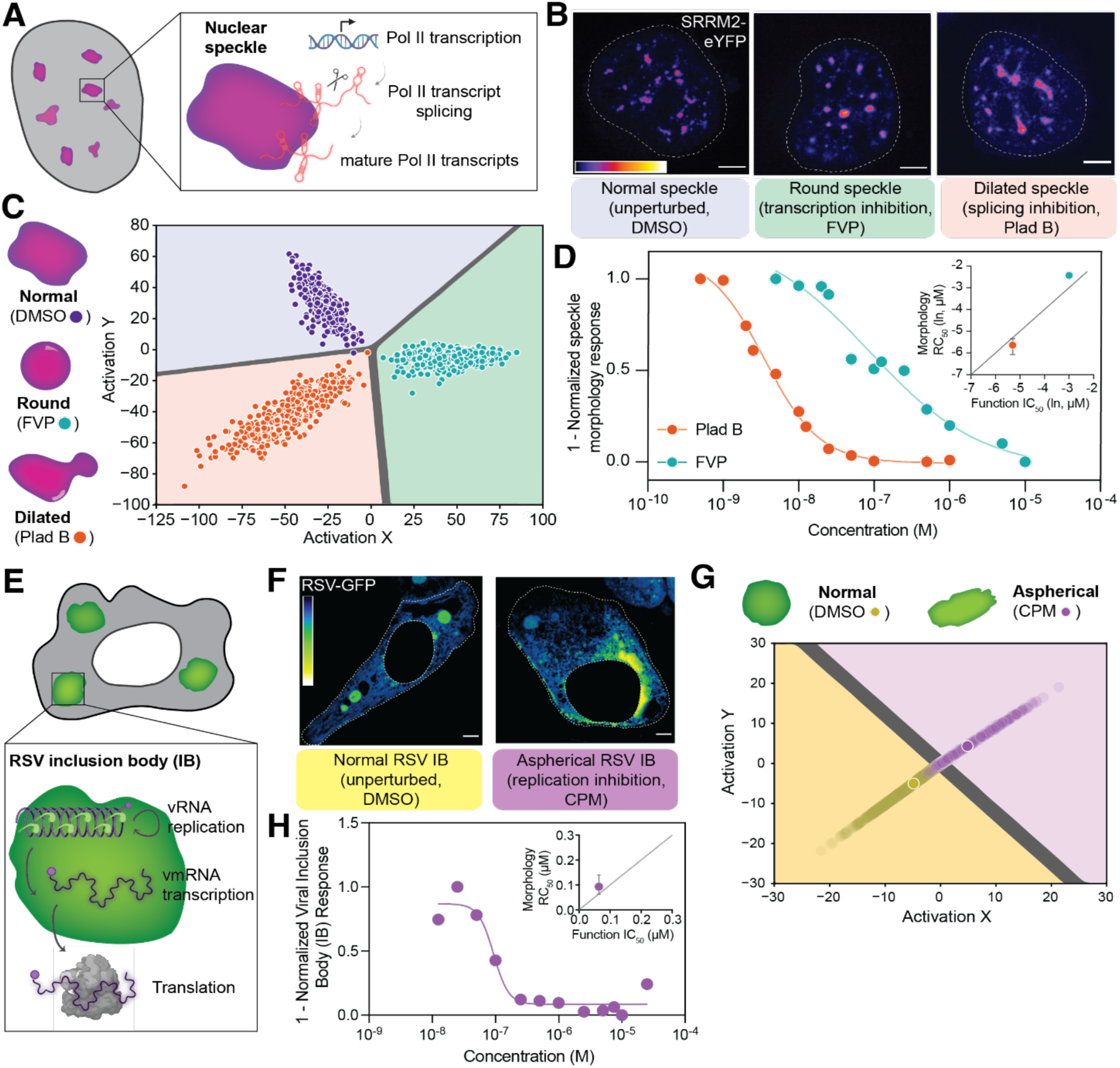
Deep-Phase applied to nuclear speckles and cytoplasmic RSV condensates quantitatively links morphology changes with inhibition of Pol II transcript biogenesis and viral RNA replication, respectively. (A) Illustration of nuclear speckle function in modulating co-transcriptional splicing of active Pol II loci. (B) Representative confocal microscopy images showing that perturbation of Pol II transcription with FVP (6.25 µM) or splicing with Pladienolide B (Plad B, 1 uM) for 2 hours alters nuclear speckle morphology compared to DMSO (0.25%) control. Speckles are visualized with endogenously tagged SRRM2-EYFP in HEK cells. Scale bar = 5 µm. (C) Activation space of the trained “speckle” neural network classifies DMSO-, FVP- and Plad B-treated cells by using the SRRM2 marker in HEK cells. Treatment time and concentration is the same as in (B). (D) Dose-response curves for FVP- or Plad B-treated cells over varying concentrations using images subjected to Deep-Phase analysis in the speckle neural network (see Figure S3A-B). Values were normalized to the lowest and highest treatment concentrations and curves were generated using a log-scaled variable slope least-squares fit. *n* ≥ 200 cells per concentration per condition. Inset shows the correlation between Deep-Phase measured RC_50_ values and literature-reported IC_50_ measurements from biochemical studies (see Table S3). In the inset, measurements were ln-transformed and a straight line was drawn through (-7,-7) for reference. Error bars for RC_50_ values represent 95% confidence intervals. (E) Illustration of human respiratory syncytial virus (RSV) cytoplasmic condensates, called inclusion bodies (IBs), that host viral RNA (vRNA) replication and viral messenger RNA (vmRNA) transcription. (F) Representative confocal microscopy images of HEp-2 cells infected with RSV-GFP for 24 hours and treated with 0.25% DMSO (left) or a replication inhibitor, cyclopamine (CPM, 10 µM), for 1 hour (right). Nuclear RSV-GFP signal was masked (see Methods). Scale bar = 5 µm. (G) Activation space of the trained cytoplasmic RSV IB neural network displaying classifications of DMSO- or CPM-treated HEp-2 cells after RSV infection. Cells were infected for 24 hours. Outlined symbols represent treatment class averages and faint dots display individual cells. (H) Dose-response curve for CPM-treated cells over varying concentrations using images subjected to Deep-Phase analysis in the cytoplasmic RSV IB neural network (see Figure S3C-D). *n* ≥ 100 cells per concentration per condition. Inset shows the correlation between Deep-Phase measured RC_50_ value and literature-reported IC_50_ measurement (see Table S4). In the inset, a straight line was drawn through (0,0) for reference and error bars for the RC_50_ value represent 95% confidence intervals.

To demonstrate the utility of Deep-Phase in measuring disruptions in RNA metabolism beyond the nucleus, we next chose to examine viral cytoplasmic condensates, which were recently identified as essential viral factories that concentrate key viral components and orchestrate viral RNA replication^83–85^. Specifically, respiratory syncytial virus (RSV) is known to form cytoplasmic condensates, referred to as inclusion bodies (IBs), that are necessary for viral RNA replication and transcription^86^ (Figure 3E). By infecting HEp-2 cells with GFP-tagged RSV, we were able to visualize round IBs throughout the cytoplasm as reported previously (Figure 3F, left). Using cyclopamine (CPM), a reported RSV replication inhibitor with activity in *in vivo* mice models, we confirmed that the morphology of RSV IBs becomes less spherical, consistent with previous reports of CPB-induced IB hardening (Figure 3F, right)^87,88^. To determine if Deep Phase can be utilized to detect these material changes, we trained a ResNet network on images of DMSO- and CPM-treated cells, focused specifically on cytoplasm (Figure S3C, see Methods). We were able to distinguish treatment morphology in the “RSV IB” activation space (Figure 3G), and then generate a dose-response curve for CPM (RC_50_ = 0.064 - 0.14 µM, 95% confidence interval, Figure 3H and S3D). This value is in close agreement with CPM-induced RSV replication IC_50_ measurement (Figure 3H inset and Table S4). These results demonstrate that Deep-Phase can serve as a broadly applicable strategy to quantitatively profile the extent of aberrant RNA condensate processes and associated biophysical changes from images of their morphology across both nuclear and cytoplasmic bodies.

### Applying Deep-Phase in a pilot drug screen identifies small molecule disruptors of nucleolar function and a unique nucleolar morphology

We next sought to utilize Deep-Phase to gain new mechanistic insights into the coupling between mesoscale condensate structure and underlying biomolecular activity, returning to the nucleolus. The nucleolus harbors RNA molecules transcribed by all three RNA polymerases^89^ as well as an abundance of RNA-binding proteins^90^, and recent studies have highlighted the role of distinct rRNA transcripts and interactions in shaping the multiphase nucleolar architecture^91–94^. We therefore reasoned that Deep-Phase could potentially uncover novel aspects of this rich RNA-centric structure-function relationship (Figure 4A).

**Figure 4.**
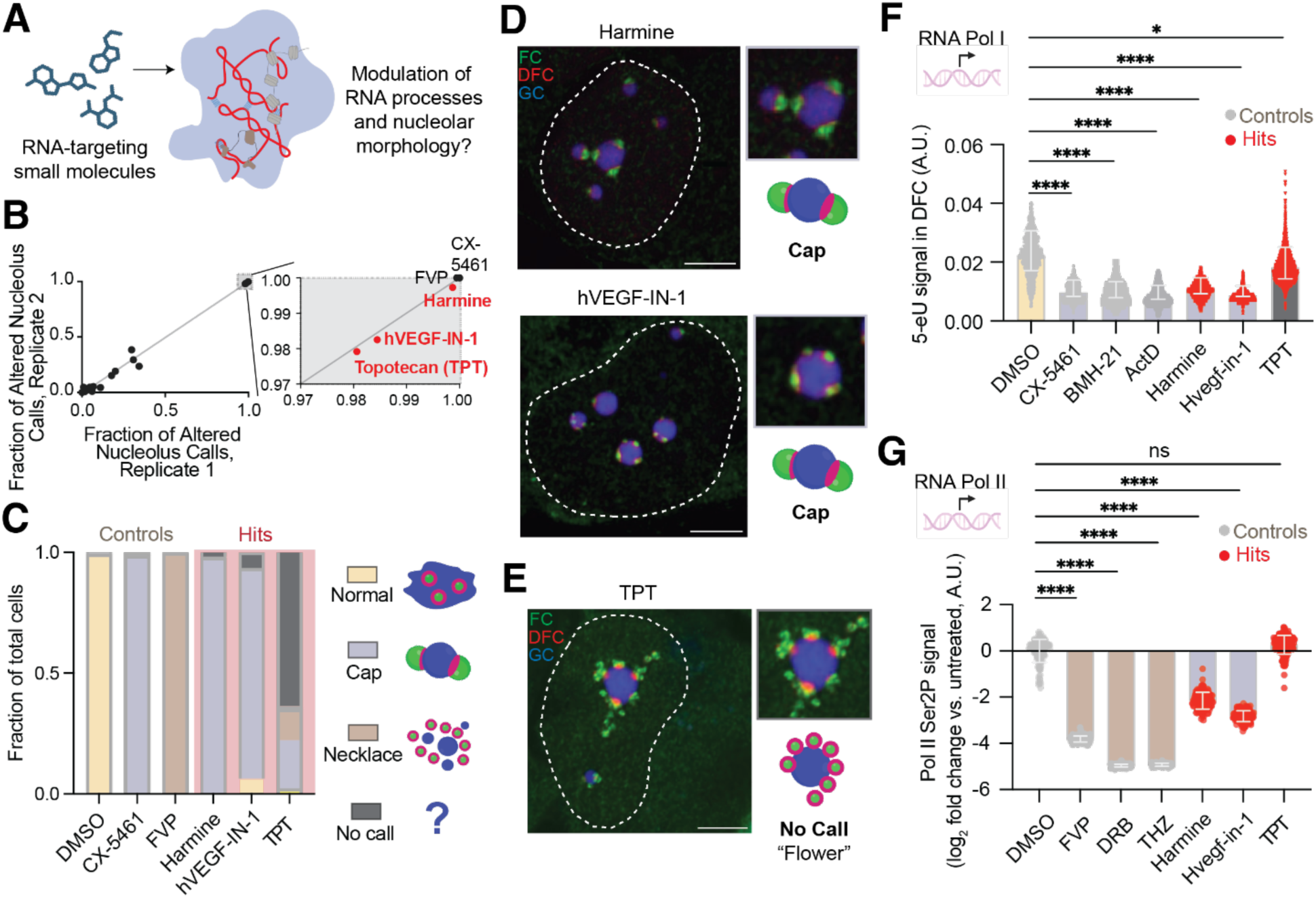
Application of Deep-Phase to a pilot drug screen of RNA-binding small molecules identifies a unique nucleolar morphology. (A) Rationale behind screening RNA-targeting ligands for nucleolar modulation. (B) Result of pilot screen showing fractions of altered nucleolar morphology calls for all treatments from two experimental replicates. Three-color nucleolar U-2 OS cells were treated with compounds for 2 hours (see Methods). Fractions were calculated by subtracting the total of cap, necklace, or no call cells as calculated by the SoftMax function from 1. A straight line was drawn through (0,0) for reference. Zoom-in box shows positive controls and hit molecules from the top right corner. *n* ≥ 240 cells per condition per replicate. (C) Combined fractions of cells from control and hit compound treatments in B from both replicates as classified by the neural network and calculated by the Softmax function (also see Figure S4A for full results). *n* ≥ 500 cells per condition. (D) Example confocal microscopy images of hits with highest “cap” calls. Scale bar = 5µm. (E) Example confocal microscopy image of a hit with the highest “No Call” calls, resembling a flower. Scale bar = 5µm. (F) Pol I transcription measurement via 5-EU metabolic labeling for controls and hit compounds. See Figure S2D for details. Bar height represents the median and error bars represent standard deviation. Statistical significance was calculated using Kruskal-Wallis test followed by Dunn’s multiple comparisons test. \**p*=0.0220, \*\*\*\**p*<0.0001. *n* = 955 (DMSO), 2772 (CX-5461), 1484 (BMH-21), 4549 (ActD), 813 (Harmine), 5067 (Hvegf-in-1), and 1871 (TPT) nucleoli. (G) Pol II transcription measurement via phosphorylated Serine 2 Pol II antibody labeling for controls and hit compounds shown as log_2_ fold changes relative to DMSO. See Figure S2E for details. Bar height represents the median and error bars represent standard deviation of raw values. Statistical significance was calculated on raw values using Kruskal-Wallis test followed by Dunn’s multiple comparisons test. *ns = not significant, \*\*\*\**p*<0.0001. *n* = 149 (DMSO), 103 (FVP), 316 (DRB), 79 (THZ), 135 (Harmine), 174 (Hvegf-in-1), and 178 (TPT) cells.

To examine this possibility, we tested a small library of RNA-binding compounds, focusing on 20 commercially available small molecules from the R-BIND 2.0 database, which contains validated RNA binders with demonstrated activity in cell culture or animal models^95^. We additionally include three known bacterial ribosome inhibitors^96^. Each molecule was screened at three different concentrations, in duplicate, and alongside cap- and necklace-forming training set controls in a 96 well plate (see Methods). To prevent false positives, we omit intrinsically fluorescent compounds by a plate reader-based quantification, which led us to discard 3 compounds (Table S5). We then plot the average frequency of altered nucleolar morphology calls for each replicate, observing a cluster of CX-5461 and FVP controls as well as three hit molecules (Harmine, hVEGF-IN-1, and Topotecan) in the upper right cluster of this graph (Figure 4B). The control treatments were correctly classified into their respective cap or necklace morphologies, while harmine and hVEGF-IN-1 were also classified as caps (Figure 4C and S4A-B), and the Topotecan (TPT) hit had a majority of “No Call” classifications (Figure 4C, S4A, and S4C). We first confirmed that hit compounds Harmine and hVEGF-IN-1 indeed caused cap morphologies, as visualized by high-resolution microscopy (Figure 4D). As expected, in follow-up functional experiments, this morphology change is accompanied by a decrease in 5-eU nucleolar signal, similar to the cap-forming positive controls (Figure 4F). However, these hit compounds also led to a slight decrease in active Pol II signal (Figure 4G), suggesting that they are non-specific intercalating transcription inhibitors, similar to Actinomycin D^97^. This is notable because both harmine and hVEGF-IN-1 are anticancer compounds whose ribosome biogenesis effects have not been reported previously^98,99^, but whose efficacy as chemotherapeutics could partly be due to inhibition of ribosome production.

The third hit compound, TPT, had the highest categorization of “No call” cells, which implies a morphology that Deep-Phase cannot classify with high confidence as normal, cap, or necklace (Figure 1C). In line with this categorization, high-resolution imaging of TPT-treated cells revealed partially inverted nucleolar morphologies, but without significant FC enlargement, that is characteristic of caps, or separation of FC and DFC layers from the GC, as is characteristic of necklaces; we named this distinct morphology as a nucleolar “flower” (Figure 4E). TPT did not cause the same functional effects as cap- and necklace-forming compounds (Figure 4F-G), suggesting that the flower morphology indeed reflects a distinct mechanism of altering nucleolar functions compared to the training set compounds.

### Characterization of the nucleolar flower morphology reveals a role of Topoisomerase I in maintaining nucleolar integrity

To gain insight into the mechanism of the TPT-induced changes in nucleolar morphology uncovered by Deep-Phase, we trained a new nucleolus network to include a set of TPT-treated cells as a distinct flower morphology category (Figure 5A), confirming that the network can classify all four morphologies with high accuracy (Figure S5A). We also ensured that the flower morphology is not a result of nucleolar protein overexpression, as wild-type U-2 OS cells stained with nucleolar antibodies displayed the same morphology after TPT treatment (Figure S5B, top). Moreover, this change is not cell type-specific, as TPT-treated HeLa cells stained with antibody markers also recapitulated the flower phenotype (Figure S5B, bottom). All phenotypes were properly classified by the updated Deep-Phase network in both cell lines (Figure S5C).

**Figure 5.**
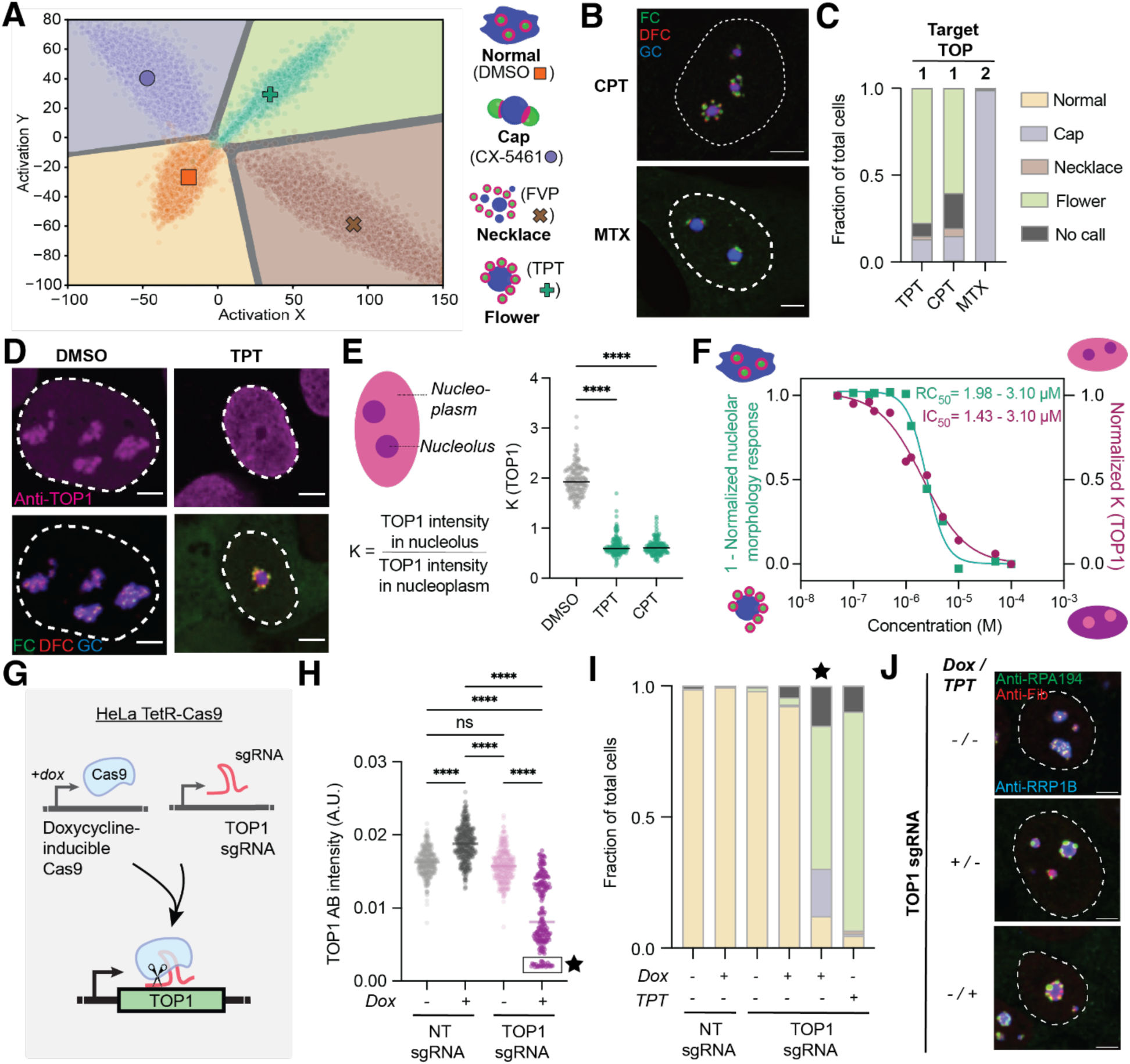
Nucleolar flower morphology is a result of TOP1 depletion. (A) Activation space of the updated neural network and schematics showing the incorporation of the nucleolar flower as a fourth nucleolar morphology class. Outlined symbols represent treatment class averages and faint dots display individual cells. (B) Representative confocal microscopy images of three-color nucleolar U-2 OS cells showing that camptothecin (CPT, 10 µM) and mitoxantrone (MTX, 1 µM) induce distinct changes in nucleolar morphology (2 hours of treatment). Scale bar = 5µm. (C) Fractions of cells from CPT and MTX treatments in B as classified by the updated neural network and calculated by the Softmax function. Topoisomerase phenotype is specific to TOP1, and not TOP2 inhibitors. *n* ≥ 100 cells per condition. (D) Representative confocal microscopy images of immunostained TOP1 in three-color nucleolar U-2 OS cells being enriched in the nucleolus (0.25% DMSO) and depleted from the nucleolus upon TPT treatment (50 µM for 2 hours). Scale bar = 5µm. (E) Quantification of change in anti-TOP1 nucleolar-nucleoplasmic partitioning upon TOP1 inhibitor treatment (50 µM TPT or 10 µM CPT for 2 hours). See Methods for details. Line is set at the median value. Statistical significance was calculated using two-tailed Mann-Whitney *U* tests. \*\*\*\**p*<0.0001. *n* = 174 (DMSO), 146 (TPT or CPT) cells. (F) Dose-response curves showing that the degree of flower phenotype formation as measured by the updated, four class Deep-Phase network (left axis) matches the extent of TOP1 nucleolar signal depletion as measured by anti-TOP1 nucleolar-nucleoplasmic partitioning (right axis). Cells were treated for 2 hours with indicated TPT concentrations. *n* ≥ 100 cells per condition. RC_50_ and IC_50_ value ranges represent 95% confidence intervals (see Table S6). (G) Schematic of HeLa TetR-Cas9 cell line generation to enable doxycycline-inducible knockout of TOP1. (H) Quantification of TOP1 antibody intensity in HeLa TetR-Cas9 cell lines with doxycycline-inducible non-targeting (NT) or TOP1-targeting (TOP1) sgRNAs. Cells were treated with 1 µg/mL of doxycycline for 6 days. Line is set at the median value. Boxed cells with the star were designated as having the highest knockout efficiency and binned for a separate Deep-Phase analysis in (I). Statistical significance was calculated using Kruskal-Wallis test followed by Dunn’s multiple comparisons test. ns = not significant, \*\*\*\**p*<0.0001. *n* = 401 (NT sgRNA - dox), 436 (NT sgRNA + dox), 313 (TOP1 sgRNA - dox), and 215 (TOP1 sgRNA + dox) cells. (I) Fractions of non-targeting (NT) or TOP1-targeting sgRNA-containing cells before and after doxycycline treatment as classified by a new, four class HeLa Cas9 network that allows for binning of SoftMax classifications based on TOP1 antibody intensity. Cells were treated with 1 µg/mL of doxycycline for 6 days or with 50 µM TPT for 2 hours. *n* ≥ 180 cells per condition, except for the binned, low anti-TOP1 intensity cells (star) that contained 33 cells. Legend is the same as in (C). (J) Representative confocal microscopy images of HeLa TetR-Cas9 cells with nucleolar marker immunostaining and containing doxycycline-inducible TOP1-targeting (sgRNAs before and after doxycycline treatment (1 µg/mL) for 6 days or after TPT treatment (50 µM) for 2 hours. Scale bar = 5µm.

We next characterized the effects of other small molecules with related modes of action. TPT was initially included in our screen as part of R-BIND based on prior reports that it disrupts pre-mRNA splicing by interfering with the interaction between NHP2L1 and U4 snRNA^100^, but is a well-established inhibitor of human DNA Topoisomerase I (TOP1)^101^. Consistent with our observed nucleolar morphology changes resulting from inhibition of TOP1, treating cells with another TOP1 inhibitor, CPT, caused a similar flower-like morphology (Figure 5B, top). However, treatment with a Topoisomerase II inhibitor, mitoxantrone (MTX)^102^, gave rise to a more cap-like morphology (Figure 5B, bottom); both morphologies were classified as expected by the updated four-class Deep-Phase network (Figure 5C). Cap-forming MTX caused a drop in Pol I transcription similar to that of cap-forming CX-5461, but substantially more than flower-forming TPT or CPT (Figure S5D). We again observed and measured a dose-dependent extent of cap or flower formation for these drugs in the updated activation space (Figure S5E-H); the MTX-induced RC_50_ for cap formation closely matched previously measured IC_50_ values for rRNA transcription inhibition (Figure S5G and Table S6).

Previous findings in other cell lines suggest that TOP1 is enriched in the nucleolus and becomes delocalized to the nucleoplasm upon TPT treatment (Figure 5D)^103–105^, suggesting that the flower morphology could result from TOP1 partitioning out of the nucleolus. We quantified this change in TOP1 partitioning across many cells, confirming that both TPT and CPT lead to decreased TOP1 partitioning in the nucleolus (Figure 5E). Treating cells with increasing TPT concentrations, we used the updated Deep-Phase network to measure an RC_50_ range of 1.98 -3.10 µM for the flower morphology, which was in very close agreement with our measurements of the IC_50_ quantified by the extent of decreased TOP1 nucleolar partitioning (1.43 - 3.10 µM, Figure 5F and S5E). The close correspondence between drug-induced TOP1 delocalization and flower morphology strongly implicates nucleolar TOP1 activity as the primary driver of this phenotype.

TPT inhibits TOP1 functioning by stabilizing the covalent TOP1-DNA cleavage complex and preventing re-ligation^101,106^. As expected for TOP1-DNA stabilization, we observed some TOP1 that remained colocalized with FC and DFC markers (Figure 5D, right), likely representing an rDNA-bound fraction of TOP1. However, this fraction was minor relative to the nucleoplasm-delocalized TOP1, suggesting that flower formation may result from a TOP1-dependent mechanism that does not involve its TPT-induced trapping on rDNA. We thus sought to functionally test whether TOP1 loss alone is sufficient to induce nucleolar reorganization, by developing a doxycycline-inducible CRISPR HeLa cell line in which we incorporated single guide (sg) RNAs targeting TOP1 (Figure 5G). After 6 days of doxycycline treatment to induce TOP1 knockout, we observed significant depletion of TOP1 protein in the cell lines containing TOP1-targeted sgRNAs (Figure 5H and S5I). TOP1 depletion was heterogeneous among these cells, which required training a modified Deep-Phase network (Figure S5J–K, also see Methods) that allowed us to use the TOP1 antibody to selectively analyze cells with the most pronounced TOP1 depletion (Figure 5H-I, star box). In this subset, cells were predominantly classified as having flower nucleoli, just as TPT-treated counterparts, whereas cells with higher residual TOP1 levels largely retained nucleolar morphologies similar to doxycycline-untreated controls (Figure 5I-J and S5M). Taken together, these results show that inhibition or depletion of TOP1 induces a distinct, cell line–independent nucleolar flower morphology arising from altered TOP1 nucleolar localization, underscoring a previously unrecognized role for TOP1 in maintaining nucleolar structure.

### TOP1 regulates rRNA processing and abundance of specific pre-rRNA intermediates

We next sought to identify what specific functional changes underlie the formation of the nucleolar flower morphology upon TOP1 inhibition. Led by the observation that TOP1-specific inhibitors induced less substantial rRNA transcription inhibition compared to TOP2 inhibitors (Figure S5D), we reasoned that the change in nucleolar morphology might be due to defects in processing of the rRNA being continually transcribed within. Indeed, CPT was previously reported to inhibit rRNA processing^48^, and the measured IC_50_ values matched the RC_50_ value we measured for the flower morphology using the updated Deep-Phase pipeline (Figure S5H and Table S6).

To test the potential role of TOP1 inhibition on rRNA processing defects, we employed 5eU-seq, which enables time-resolved measurements of rRNA processing in the nucleolus^94^. Briefly, cells were first treated with TPT or DMSO for 1 hour, followed by a 15-minute pulse of 5-ethynyl uridine (5-eU) to label nascent transcripts, and then chased for varying time intervals up to 90 minutes (Figure 6A and S6A). 5eU-seq analysis revealed that TPT treatment causes significant rRNA processing defects across multiple internal cleavage sites of the 47S pre-rRNA. Interestingly, the most pronounced defects occur at middle and late ribosome assembly steps within the nucleolus (sites 1, 2, and 3’), while co-transcriptional processing at the 5’ ETS (site 01) remains largely unaffected (Figure 6B). However, these processing defects mirrored those recently reported upon FVP treatment^94^, which results in a qualitatively different morphology - the necklace - that can be clearly distinguished by Deep-Phase (Figure 1B). This raises a key question: how can similar processing defects give rise to morphologically distinct nucleoli?

**Figure 6.**
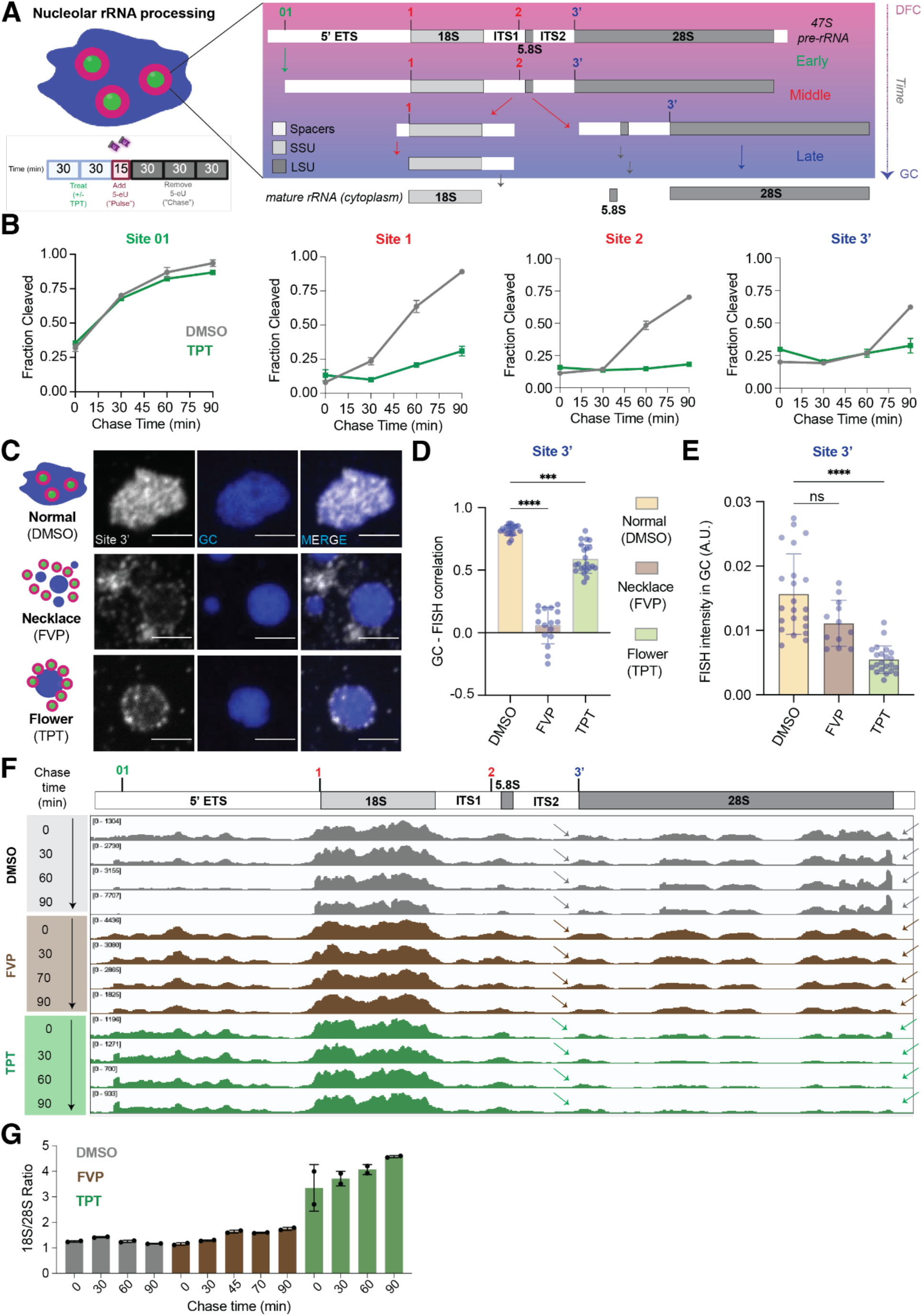
TPT and FVP treatments differentially affect localization and abundance of late-stage pre-rRNA intermediates. (A) Simplified schematic of nucleolar pre-rRNA processing steps highlighting early (green), middle (red), and late (blue) cleavage events over time that can be captured with the 5eU-seq labeling paradigm^94^. DFC = dense fibrillar component, GC = granular center, SSU = small subunit, LSU = large subunit. ( B) Results of 5eU-seq experiment showing that TPT (50 µM) induces global rRNA processing defects as assessed by pulldown and sequencing. See Figure S6A and Methods for details. Data is shown as mean ± S.E.M. *n* = 2 replicates per time point. (C) Representative confocal microscopy images of pre-rRNA junction 3’ by RNA-FISH reveals differential localization patterns for FVP-(1 µM, 2 hours) versus TPT-treated (50 µM, 2 hours) three-color nucleolar U-2 OS cells, despite similarities in rRNA processing defects. Scale bar = 2.5 µm. (D) Quantification of RNA FISH signal correlation with GC for site 3’ from images shown in Figure 6C (see Methods). Bar height represents the median and error bars represent standard deviation. Statistical significance was calculated using Kruskal-Wallis test followed by Dunn’s multiple comparisons test. \*\*\**p* = 0.0004, \*\*\*\**p*<0.0001. *n* = 20 (DMSO), 16 (FVP), 22 (TPT) cells. (E) Quantification of RNA FISH signal intensity in the GC for site 3’ from images shown in Figure 6C (see Methods). Bar height represents the median and error bars represent standard deviation. Statistical significance was calculated as in (D). \*\*\*\**p*<0.0001, *ns* = not significant. *n* is the same as in (D). (F) Drop in coverage of 5eU-seq reads in the second half of the rRNA transcript in TPT-treated compared to DMSO-treated or FVP-treated cells (arrows). *n* = 2 replicates per time point, see Figure S6B for the second replicate and Figure S6C for DMSO control from the FVP experiment. FVP data was previously reported^94^. (G) Quantification of 18S (first half, SSU) over 28S (second half, LSU) 5eU-seq reads regions after DMSO, FVP, or TPT treatment at different chase timepoints displays decrease in 28S reads for TPT treatment. Data is shown as mean ± S.E.M. *n* = 2 replicates per time point; FVP data was previously reported^94^.

The most conspicuous structural difference between flower and necklace morphologies is the presence of a defined DFC–GC interface in flowers, and its apparent loss in necklaces, where GC droplets largely detach from the DFC. Previous work has suggested that late-stage rRNA junctions whose cleavage in the GC is essential to formation of the ribosomal large subunit (LSU) are misprocessed during FVP treatment, and improperly accumulate in the nucleoplasm between the DFC and GC^94^. This prompted us to examine whether the spatial distribution of misprocessed LSU species differs between FVP and TPT treatment conditions. Using RNA FISH, we found that upon TPT treatment, pre-rRNAs containing the late-stage site 3’ were localized entirely within the GC, similar to the DMSO-treated control, whereas FVP-treated cells showed pre-rRNAs localized predominantly in the nucleoplasm, quantified as a substantially lower GC-FISH signal correlation (Figure 6C-D). We additionally noted and quantified that the overall site 3’ FISH signal in the GC was lower in the TPT-treated compared to control and FVP-treated cases (Figure 6C and 6E). We therefore reasoned that the difference in the relative abundance of misprocessed late-stage, LSU-forming rRNA intermediates between FVP and TPT treatments could be responsible for the formation of distinct nucleolar morphologies. Consistent with this, 5eU-seq read coverage plots revealed a striking reduction in reads mapping to the LSU-containing, 3’ half of the 45S transcript following TPT but not FVP treatment (Figure 6F and S6B-C). We quantified this difference in the form of 18S/28S ratios as a proxy for SSU vs. LSU reads over time (Figure 6G). This difference likely results from incomplete transcription through the 3’ end of rDNA due to unresolved supercoiling normally relieved by TOP1^107^, or possibly through selective degradation of site 3’-containing LSU precursors.

We next sought to assess how pre-rRNA abundance and localization changes under prolonged TOP1 loss rather than short inhibition (hours) via TPT. 5eU seq was not possible in the TOP1 knockout system due to heterogeneous knockout efficiency (Figure 5H), but we were able to perform RNA FISH experiments in these cells. Strikingly, in flower-morphology cells with low TOP1 signal, we observed a near-complete loss of FISH signal for site 3’ relative to non-targeting sgRNA cells (Figure S6D-E). This absence of site 3′ signal is consistent with the nucleolar flower resulting from cells chronically failing to generate sufficient 28S-containing pre-rRNAs to maintain normal structure. The retained site 3’-containing transcripts detected in the GC of TPT-treated cells (Figure 6C and 6E) therefore likely reflect rRNAs that localized to the GC prior to drug treatment. Together, these findings all point to the differential accumulation of misprocessed rRNA precursors at the DFC–GC interface as driving the distinct necklace versus flower morphologies.

### Nucleolar DFC-GC interface maintenance depends on the identity and abundance of rRNA precursors

The model that an abundance of misprocessed rRNA compromises the integrity of the DFC-GC interface predicts that reducing the overall load of misprocessed transcripts in FVP-treated cells should restore the DFC–GC boundary; given the otherwise similar patterns of rRNA misprocessing between FVP-treatment and TOP1 disruption, this could even potentially recapitulate the flower morphology. To this end, we designed a two-step perturbation experiment: we first treated cells with FVP to induce global rRNA processing defects and disrupt the DFC–GC interface, and then added CX-5461 to suppress rRNA transcription and thereby reduce the accumulation of misprocessed intermediates (Figure 7A). Strikingly, this sequential treatment produced a nucleolar morphology that closely resembled the flower phenotype, and was classified as such by the updated Deep-Phase pipeline (Figure 7B–C). As expected, FISH experiments for site 3’ revealed increased FISH-GC correlation as well as a concomitant decrease in FISH signal intensity for the sequential treatment case compared to FVP only treatment (Figure 6SF-H). These findings suggest that reducing the concentration of misprocessed rRNA intermediates allows the DFC–GC interface to be re-established, consistent with a model in which proper organization of nucleolar phases depends on continuous transcription and processing of rRNA, with differential changes leading to cap, necklace, or the TOP-1-associated flower morphologies (Figure 7D).

**Figure 7.**
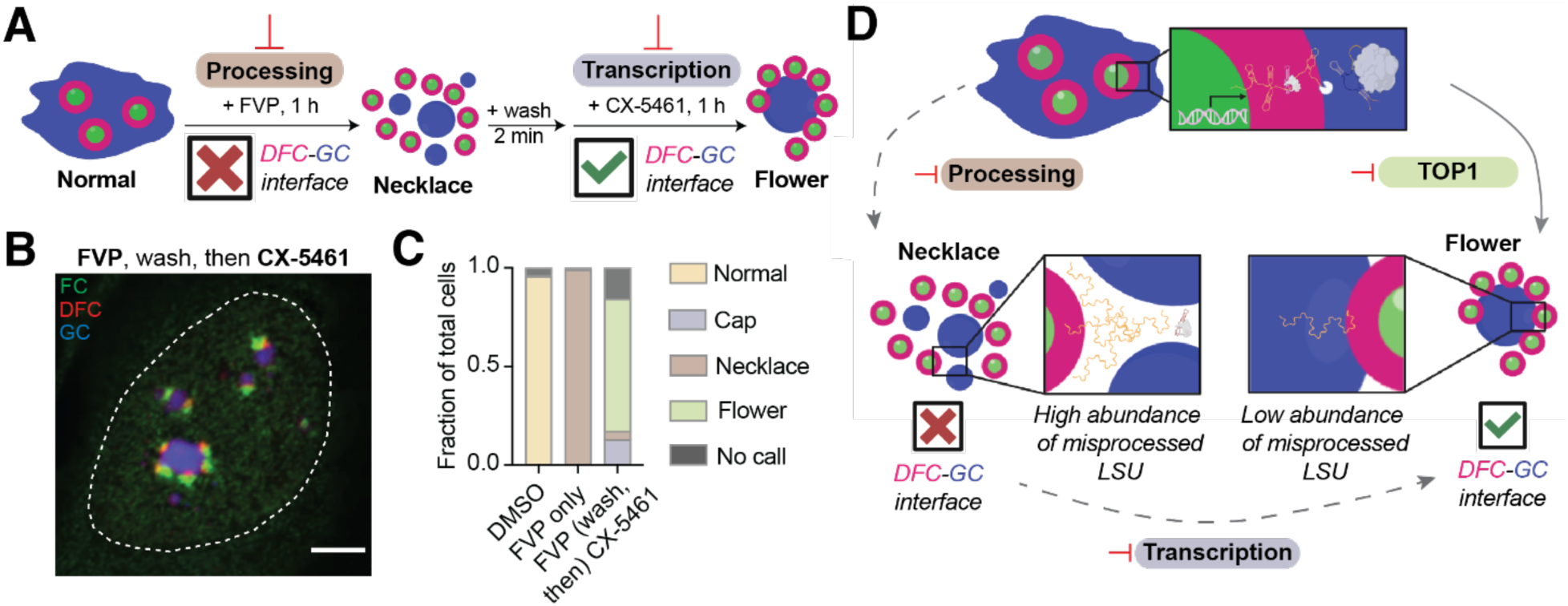
Sequential processing and transcriptional inhibition recapitulate the flower morphology. (A) Hypothesis and experimental schematic for modulating the DFC-GC interface presence by inhibiting processing with FVP (1 µM, 1 hour) followed by a wash (2 minutes), and then transcription inhibition with CX-5461 (1 µM, 1 hour). (B) Representative confocal microscopy image of experiment described in A. Scale bar = 5µm. (C) Deep-Phase output of individual cells as classified by the updated four-class neural network and calculated by the Softmax function. *n* ≥ 70 cells. (D) Proposed model for rRNA identity and abundance-dependent boundary of nucleolar subphases. In a normal nucleolus with proper rRNA processing, the FC, DFC, and GC layers are organized concentrically. Upon processing inhibition (e.g. FVP treatment, left dashed arrow), the concentric layering is disrupted when the nucleolus forms the necklace morphology, with misprocessed LSU rRNAs accumulating between the DFC (red) and GC (blue) interface. If transcription is subsequently inhibited (e.g. CX-5461 treatment, bottom dashed arrow), the abundance of misprocessed rRNA LSU intermediates is lowered, resulting in the re-establishment of the DFC-GC interface such as that observed with the flower morphology. Instead of this two-step process, the same flower morphology can form when TOP1 is depleted from the nucleolus, either chemically (e.g. TPT treatment) or genetically (e.g. CRISPR knockout).

## DISCUSSION

A long-standing challenge in cell biology is connecting mesoscale structural organization with underlying molecular driving forces. This challenge is particularly well defined, but still poorly understood, in biomolecular condensates, whose emergent phase behavior is a direct result of the thermodynamics and biochemical reactions occurring within. Here, we develop a deep learning-based approach - Deep-Phase - that enables quantitative study of condensate structure-function relationships and their disease-relevant implications. By measuring dose-response and kinetic changes in nucleolar, nuclear speckle, and viral condensate morphology, Deep-Phase provides a platform for unbiased inference of molecular-level activities from mesoscale condensate structures. The quantitative power of Deep-Phase is underscored by the close correspondence between our measured RC₅₀ values for drug-induced condensate morphology changes and IC₅₀ values from underlying functional perturbations, across distinct assay systems and cell types. This sensitivity allows morphological phenotypes to serve as direct proxies for target engagement, even in diverse cellular backgrounds.

Applying the Deep-Phase framework in a small molecule screen for nucleolar morphology changes demonstrates its potential as a high content platform for phenotypic screens, where most novel biological interactions and first-in-class compounds have been discovered^108^. Deep Phase can uncover novel nucleolar morphological changes that reveal new aspects of nucleolar biology, as demonstrated by our finding of a previously uncharacterized nucleolar “flower” morphology resulting from TOP1 inhibition. Moreover, we demonstrated that Deep-Phase can be extended to detect RNA-dependent changes in other condensates, including viral condensates and nuclear speckles. With the growing recognition of RNA as a therapeutic target, this is particularly relevant for the rapidly expanding field of RNA-targeting small molecules^109–111^; we envision that our imaging-based approach could offer a powerful, cell-based screening platform for discovering new therapeutic modalities that engage condensate-localizing RNA and RNA-binding proteins.

Our small molecule screen led to the discovery of an unexpected function for TOP1 in ribosome biogenesis: its inhibition causes a flower morphology arising from distinct pre-rRNA processing defects, including reduced transcription and processing of LSU precursors. These findings reveal that in addition to TOP1’s established role in resolving DNA supercoiling - particularly on active, highly transcribed genes such as rDNA repeats^112,113^ - TOP1 also plays a previously unrecognized role in coordinating rRNA processing, likely through its recently described non-canonical impacts on RNA metabolism, including direct binding and cleavage of RNA.^114,115^ Loss of TOP1 function is thus similar to combined misprocessing and depletion of accumulating rRNA intermediates, as we demonstrated with sequential FVP and CX-5461 treatment, which recapitulates the flower morphology as a mesoscale manifestation of these coupled molecular-level defects. Notably, this function is unique to TOP1, as TOP2 inhibition instead leads to the canonical cap morphology, distinguishable by Deep-Phase. Taken together, these findings demonstrate that nucleolar morphology changes detected by Deep-Phase are a quantitative and mechanistic fingerprint of the rRNA processing state, shaped by the interplay between RNA concentration, phase boundaries, and biochemical function.

We envision a number of future applications of Deep-Phase to answer pressing questions in condensate biology. For example, the ability to distinguish nucleolar responses to inhibition of TOP1 and TOP2—related enzymes from topoisomerase subclasses—illustrates its potential for resolving functional phenotypes across structurally similar targets. This method could therefore be leveraged to systematically probe genetic determinants of condensate form and function, providing insights into how these bodies are shaped by cellular pathways and mutations. Such a framework could uniquely enable the investigation of condensate functions of disordered or transiently interacting proteins, which often evade detection by structure-based approaches, and may uncover previously unrecognized regulators of condensate organization. Its ability to detect subtle structural perturbations also makes Deep-Phase well-suited for studying disease states where aberrant condensate morphologies may indicate altered functional pathways or molecular species imbalances. We propose that Deep-Phase could provide a scalable approach for functional phenotyping of cancer cells, with potential applications in drug screening, mechanism-of-action studies, identification of morphology-based biomarkers, and drug response heterogeneity detection at single-cell resolution.

More broadly, Deep-Phase offers a generalizable strategy for decoding the functional logic of condensate organization from images alone. By enabling quantitative, high-throughput inference of biochemical states from mesoscale structures, it opens the door to functional condensate profiling at scale - across cell types, perturbations, and disease contexts. Given the increasing recognition of condensates as ubiquitous hubs of biological activity throughout the cell, Deep-Phase provides a platform for uncovering new principles of intracellular organization, and for translating condensate morphology into a readout of cellular state with diagnostic and therapeutic potential.

### Limitations of the study

While Deep-Phase provides a powerful platform for quantitatively linking condensate morphology to molecular function, several limitations remain. First, the approach primarily establishes correlations between structural and functional changes, and further experiments are needed to yield full mechanistic insight. Second, although the method captures mesoscale organizational changes, finer molecular-scale rearrangements within condensates may be missed, and changes in smaller condensates that are below the resolution limit of high-content imaging equipment may be difficult to capture. Integrating correlative imaging and adaptive training with super-resolution or simulation images^116,117^ will help address these challenges. Alternatively, advances in molecular-resolution, activity-based, and engineered repeat operon or optogenetic imaging readouts may be employed^118,119^. Third, like many deep learning-based methods, the underlying neural networks function as “black boxes,” making it difficult to definitively interpret the specific image features driving classification decisions. Applying techniques that explain deep learning classifier predictions^120^ will further improve the interpretability of this framework. Finally, a limitation of the current framework is that it relies on starting with condensates whose molecular functions and perturbation mechanisms are already partially understood, as the ability to assign functional meaning to structural changes depends on having known benchmarks, such as drugs with defined effects on specific biochemical pathways. This makes Deep-Phase particularly powerful for hypothesis-driven studies and mechanistic dissection, but potentially less straightforward to apply to novel, poorly characterized condensates without complementary functional assays. Nevertheless, as structural signatures are mapped across broader perturbation spaces, Deep-Phase may ultimately enable *de novo* functional inference from morphology alone, even in the context of novel condensate types.

## Resource availability

### Lead contact

Requests for further information should be directed to the lead contact, Clifford P. Brangwynne.

### Materials availability

All materials and resource requests should be directed to and will be fulfilled by the lead contact, Clifford P. Brangwynne.

### Data and code availability

Data in this manuscript will be shared by the lead contact upon request. All image analysis pipelines, including custom scripts and Deep-Phase software with example images for testing, are available at https://github.com/SoftLivingMatter/Deep-Phase with the published version maintained at https://doi.org/10.5281/zenodo.16103290.

## Acknowledgments

We are grateful to Dr. Hahn Kim and Prof. Mengdi Wang for helpful early discussions regarding this study; Evangelos Gatzogiannis for help with automated microscopy protocol setup; Christina DeCoste for assistance with cell sorting; David Sanders for plasmids; Amy Strom and Lindsay Becker for thoughtful feedback on the manuscript, and all Brangwynne lab members for helpful discussions as well as experimental support. We are additionally pleased to acknowledge that high-performance computing work in this manuscript, including all Deep-Phase training and analyses, was performed using the the Princeton Research Computing resources at Princeton University, a consortium of groups led by the Princeton Institute for Computational Science and Engineering (PICSciE) and Office of Information Technology’s Research Computing. This work was supported by the Howard Hughes Medical Institute (HHMI), the AFOSR MURI (FA9550-20-1-0241), the Princeton Center for Complex Materials, a MRSEC (NSF DMR-2011750), the St. Jude Collaborative on Membraneless Organelles, and the Chan Zuckerberg Initiative Exploratory Cell Network. S.A.Q. is supported by the HHMI Hanna H. Gray fellowship. K. Antunes Fernandes received the Pew Latin American Fellowship, funded by The Pew Charitable Trust, and the Princeton Molecular Biology Postdoctoral Research Fellowship. AI Lim is supported by Searle Scholar Program and The Branco Weiss Fellowship. L.W.W. is supported by the NSF GRFP.

## AUTHOR CONTRIBUTIONS

A.D. and C.P.B. designed the study. A.D., S.A.Q., N.J-L., K.H.A., L.J., and L.W.W. performed experiments with advice from A.I.L. and C.P.B. T.J.C. trained and optimized all deep neural networks. A.D., T.J.C., and S.A.Q. performed quantitative imaging and genomics data analysis. A.D. and C.P.B. wrote the manuscript with input from all authors. A.D. made the figures with contributions from all authors.

## DECLARATION OF GENERATIVE AI AND AI-ASSISTED TECHNOLOGIES

During the preparation of this work, the authors used chatGPT in order to improve readability and clarity of some written sections. After using this tool, the authors reviewed and edited the content as needed and take full responsibility for the content of this publication.

## SUPPLEMENTAL FIGURES

**Figure S1.**
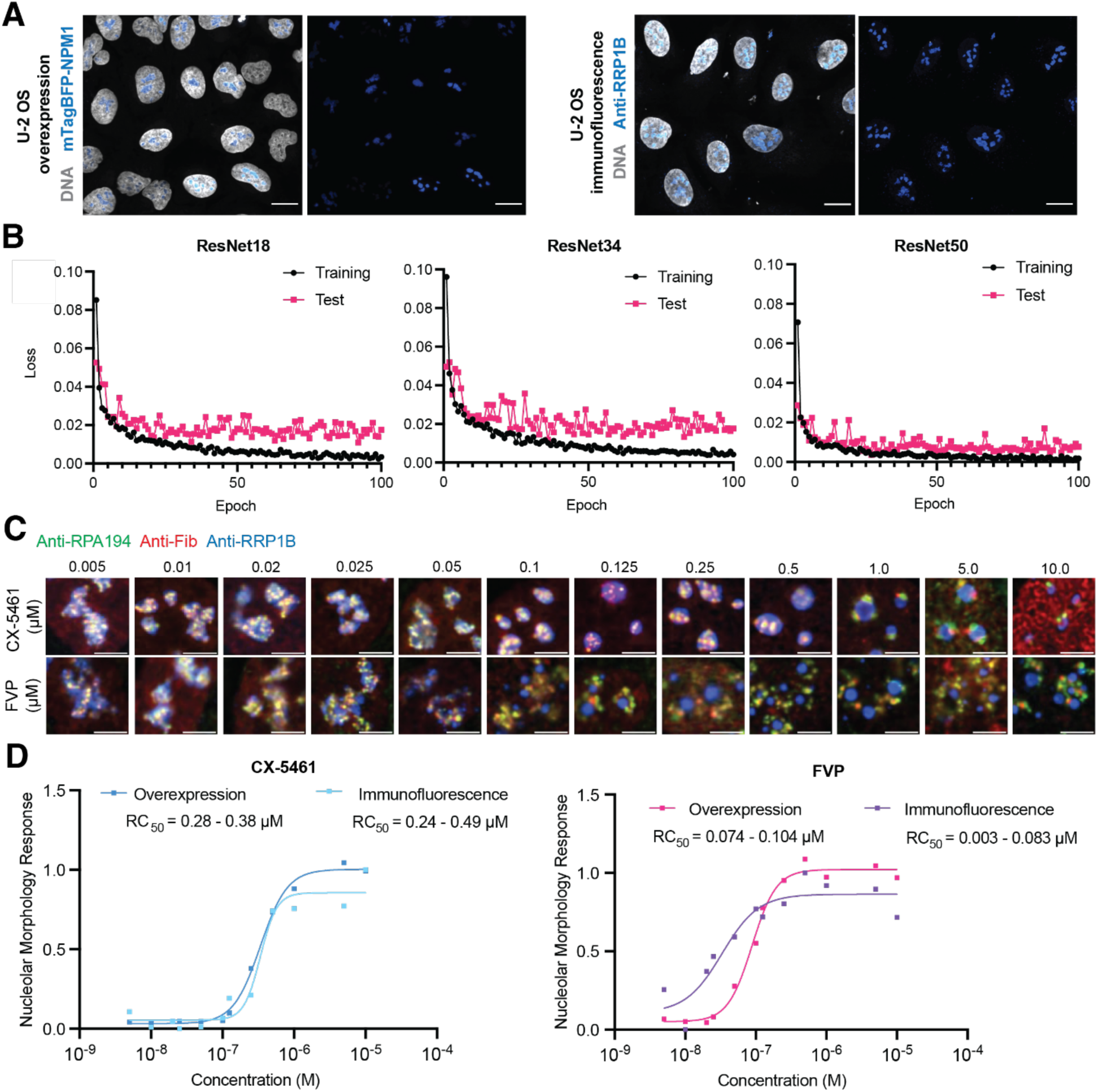
Validation and optimization of Deep-Phase methodology in the three-color U-2 OS overexpression line and ResNet networks that detect changes in nucleolar morphology, related to Figures 1-2. (A) Representative confocal microscopy images comparing nucleolar morphology in the three-color U-2 OS nucleolar line overexpressing RPA16-GFP (FC marker, not shown), Nop56-mCherry (DFC marker, not shown), and mTagBFP2-NPM1 (GC marker) with wild-type U-2 OS cells stained with Anti-RRP1B (GC marker). DNA was stained with SiR-DNA. Scale bar = 10 µm. (B) Cross entropy loss of correct image classification (normal, cap and necklace) during network training by three ResNet architectures for training and test sets. Networks differ in the number of convolutional layers (18, 34, and 50). (C) Representative confocal microscopy images of antibody-stained wild-type U-2 OS cells treated with increasing concentrations of CX-5461 (top row) and FVP (bottom row) for 2 hours. Scale bar = 5 µm. (D) Dose-response curves and RC_50_ values of antibody-stained wild-type U-2 OS cells (“Immunofluorescence”) compared to the originally used three-color U-2 OS cell line overexpressing nucleolar compartment markers (“Overexpression”) after treatment with increasing concentrations of CX-5461 and FVP for 2 hours. *n* ≥ 55 per condition.

**Figure S2.**
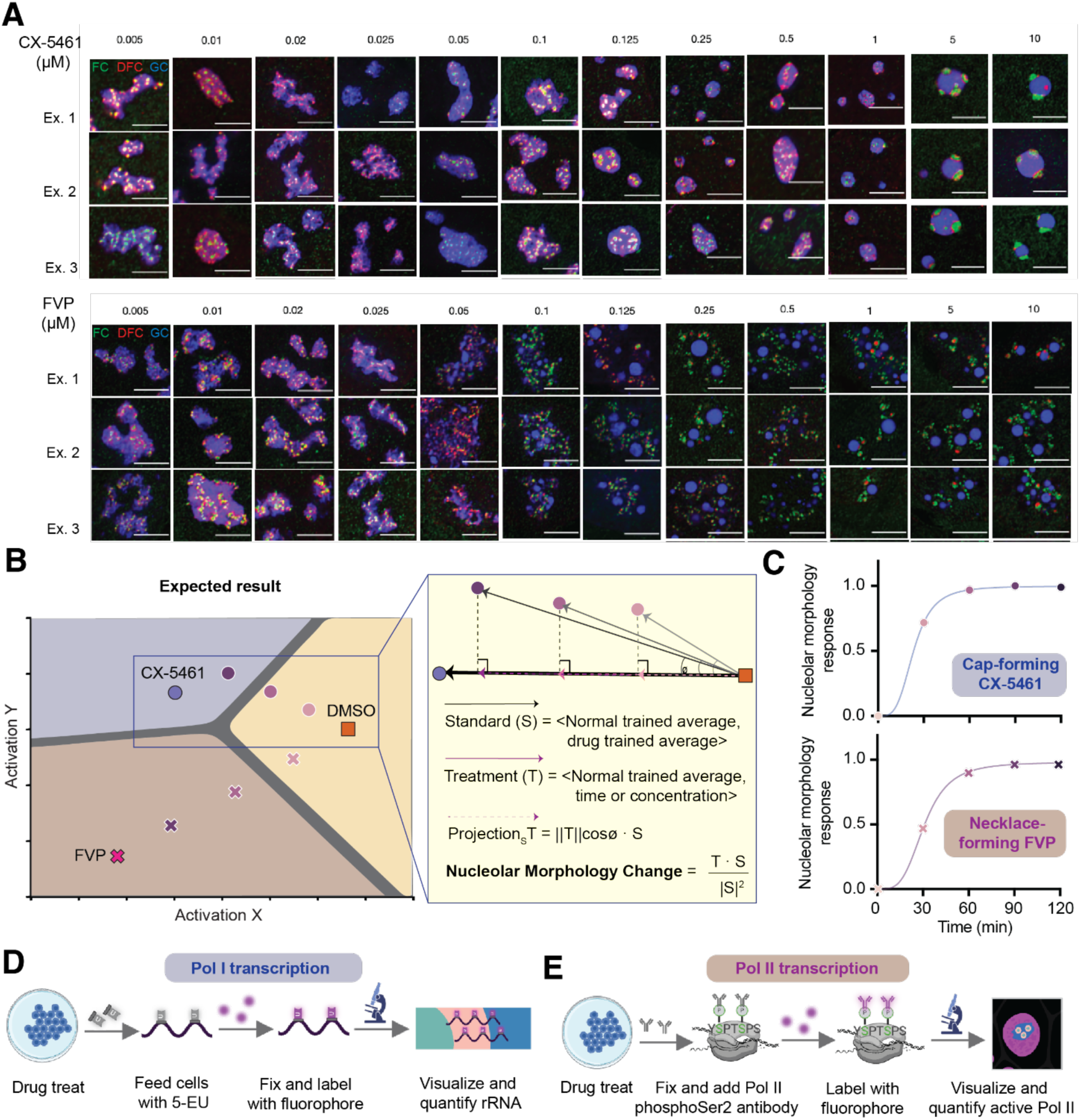
Quantifying gradual changes in nucleolar morphology by Deep-Phase after time- and dose-dependent drug treatments and linking them to functional perturbations, related to Figure 2. (A) Examples of nucleoli from different cells treated with varying CX-5461 (top) or FVP (bottom) concentrations for two hours. Scale bar = 5µm. (B) Vector algebra used to calculate a nucleolar morphology response in the two-dimensional activation space. See Figure 2C-D for time- and concentration-dependent morphology changes upon drug treatment, respectively. See Table S1-S2 for RC_50_ measurements. (C) Nucleolar morphology change upon treatment with CX-5461 (10 µM, top) or FVP (6.25 µM, bottom) in 30 minute intervals. The change was quantified using the vector algebra as described in Figure S2B from the activation space shown in Figure 2C. *n* ≥ 218 cells per condition. (D) Functional readout assessing levels of RNA Pol I transcription for cap-forming drugs (see Methods). During the last 15 minutes of drug treatment, cells are fed 5-ethynyl-uridine (5-eU) that can be visualized with a click-chemistry-containing fluorophore after fixation (see Methods). 5-eU signal intensity in the DFC is quantified as a proxy of rRNA levels. (E) Functional readout assessing levels of RNA Pol II transcription for necklace-forming drugs. Following drug treatment and fixation, cells are immunolabeled with an antibody detecting active Pol II transcription (see Methods). Nuclear antibody signal is quantified as a proxy of Pol II CDK9-dependent Serine 2 phosphorylation.

**Figure S3.**
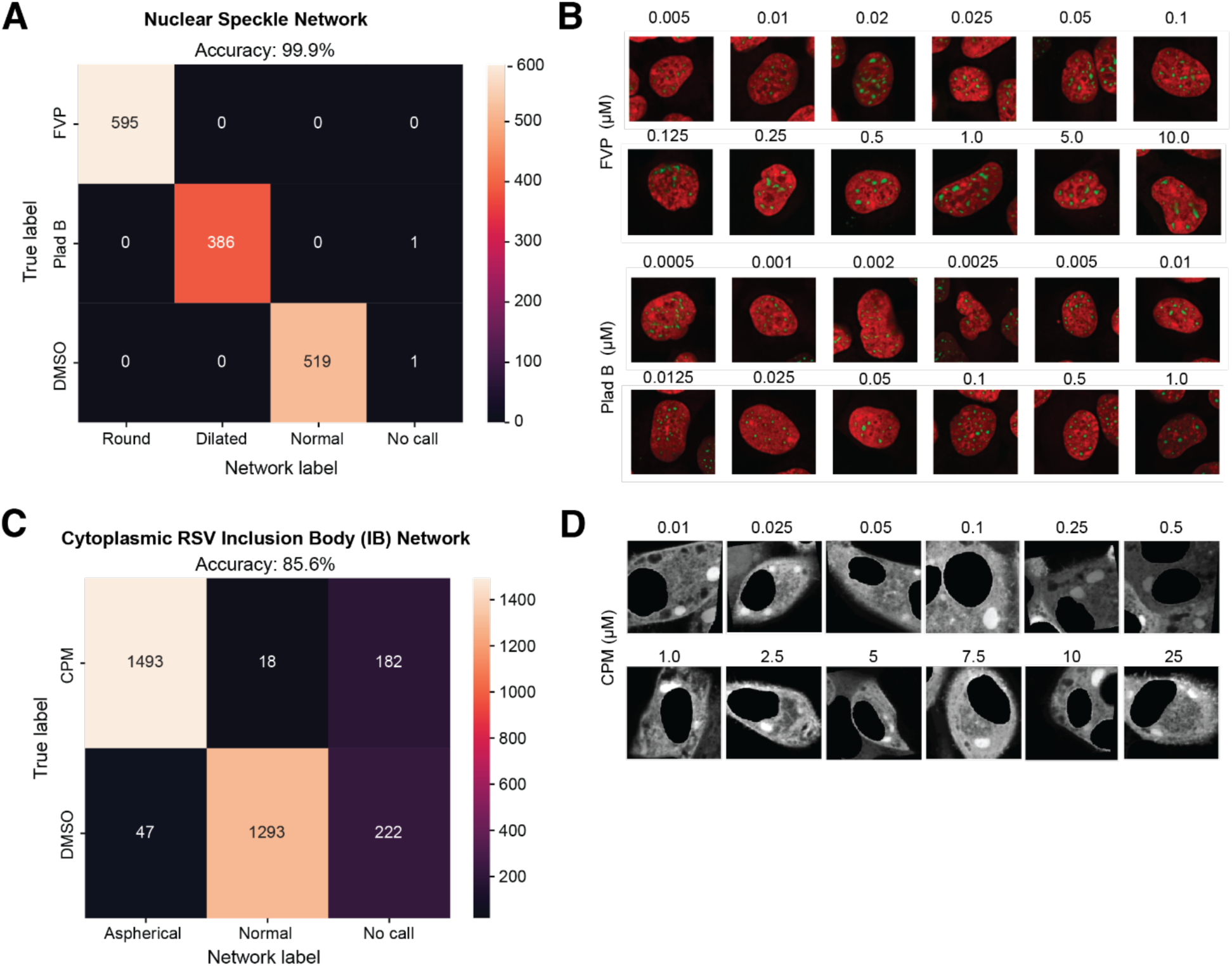
Extension of the Deep-Phase approach to detect drug-induced disruptions in nuclear speckle and viral cytoplasmic condensate processes, related to Figure 3. (A) Confusion matrix for the nuclear speckle network trained with images of SRRM2 HEK cells from the test set displays high accuracy for all three speckle morphology classes (see Methods). Cells were treated with FVP as a Pol II transcriptional inhibitor (6.25 µM), or Pladienolide B (Plad B, 1 µM) as a splicing inhibitor for 2 hours. (B) Example images of cropped HEK cells from the speckle network endogenously tagged with a nuclear speckle marker EYFP-SRRM2 (green) and stained with SiR-DNA. Cells were treated with increasing concentrations of FVP or Plad B for 2 hours to generate dose-response curves and RC_50_ values. See Table S3. (C) Confusion matrix for the cytoplasmic RSV inclusion body network trained with images of HEp-2 cells after infection with RSV-GFP and treatment with viral replication inhibitor, CPM (10 µM), for 1 hour. Nuclei were segmented out to ensure that only cytoplasmic RSV signal was used in the network training and classification (see Methods). (D) Example images of cropped RSV-infected HEp-2 cells treated with increasing concentrations of CPM for 1 hour used to generate dose-response curves and RC_50_ values. See Table S4.

**Figure S4.**
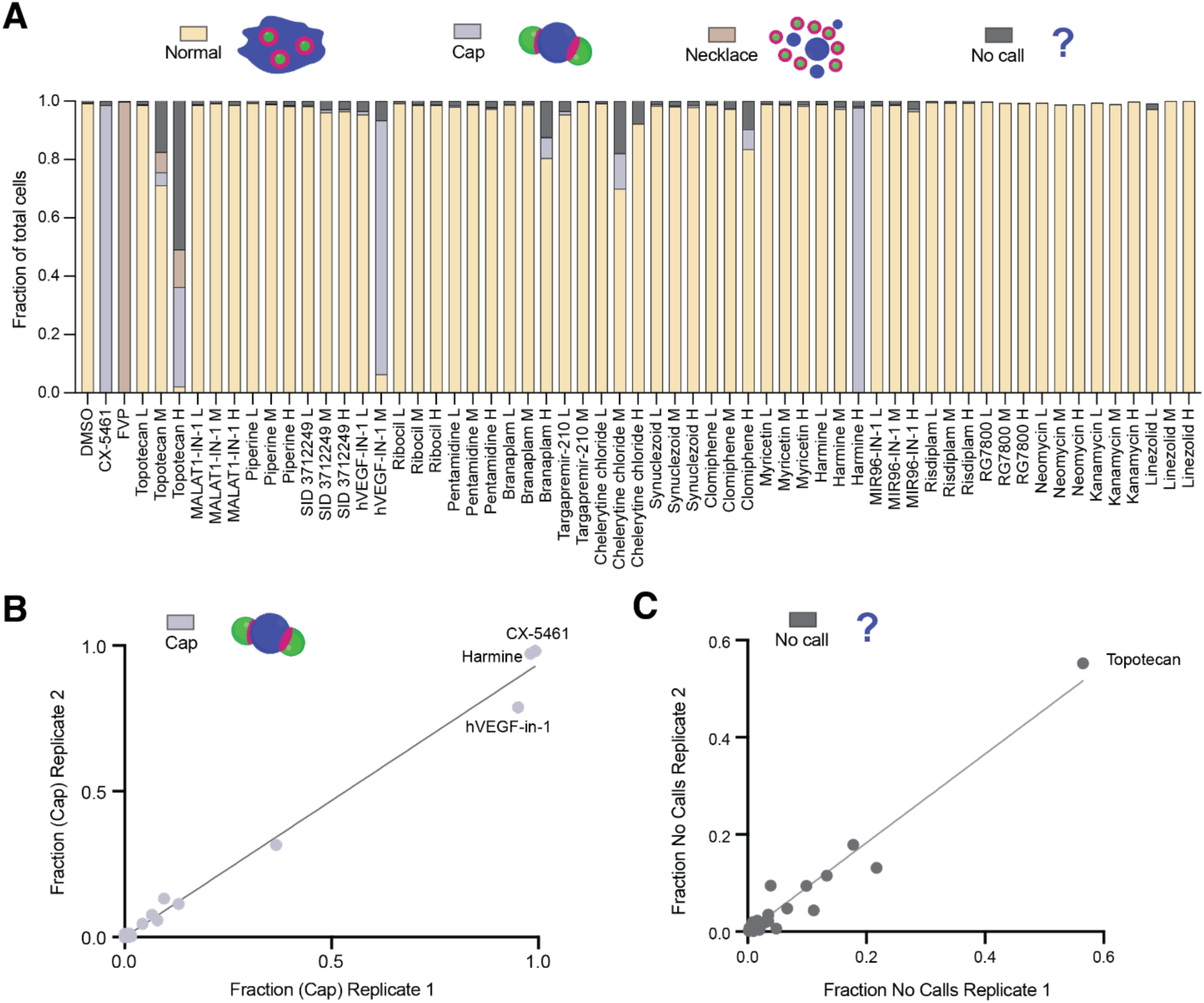
Results of Deep-Phase application in a proof-of-concept screen of RNA-targeting small molecules, related to Figure 4. (A) Fractions of total cells classified as having a normal, cap, necklace, or “no call” nucleolar morphology in the pilot screen of RNA-targeting molecules; also see Methods. Fractions were generated by feeding images from two experimental replicates to the nucleolar Deep-Phase network and generating class probabilities with the SoftMax function. L = low concentration (1 µM), M = medium concentration (10 µM), H = high concentration (50 µM). *n* ≥ 247 cells. (B) Fractions of cells with “cap” morphology class as generated by Deep-Phase and calculated by the SoftMax function from two experimental replicates. (C) Fractions of cells with “No Call” morphology class as generated by Deep-Phase and calculated by the SoftMax function from two experimental replicates.

**Figure S5.**
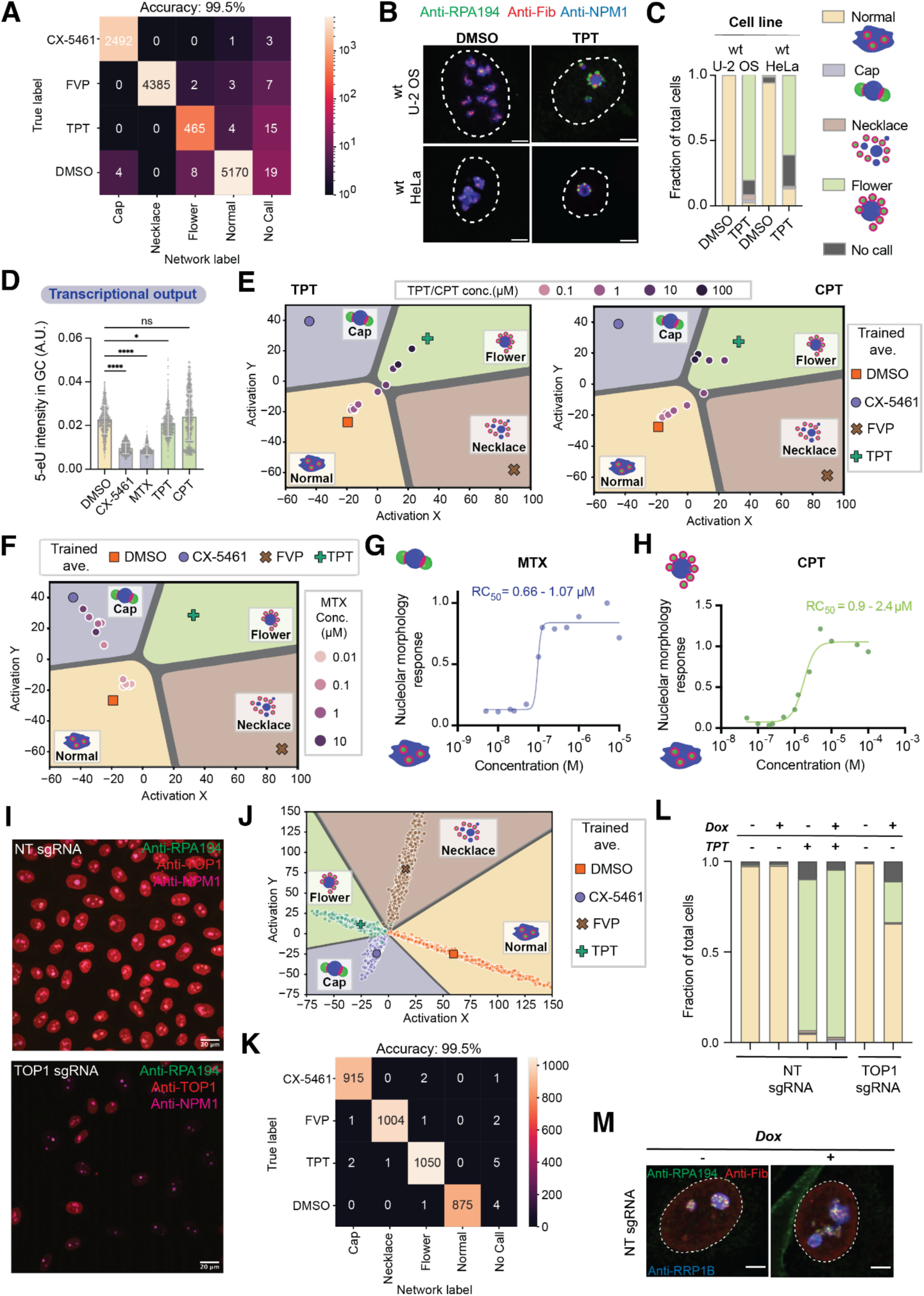
Updated Deep-Phase nucleolar network classifies and quantifies distinct flower morphology formation upon TOP1 nucleolar depletion with chemical or genetic perturbations, related to Figure 5. (A) Confusion matrix for the updated nucleolar Deep-Phase network that contains TPT-treated cells as the fourth nucleolar morphology class (flower). The three-color nucleolar U-2 OS cell line was treated with 50 µM of topotecan (TPT) for 2 hours. (B) Representative confocal microscopy images of DMSO-(0.25 %) or TPT-treated (50 µM, 2 hours) wild-type U-2 OS and HeLa cells after immunostaining with FC (RPA-194), DFC (Fibrillarin), and GC (NPM1) markers. Scale bar = 5 µm. (C) Classifications of DMSO- or TPT-treated wild-type U-2 OS and HeLa cells by the new, four-class nucleolar Deep-Phase network. Fractions were calculated from SoftMax function outputs. *n* ≥ 350 cells (U-2 OS) and ≥ 800 cells (HeLa). (D) Quantification of Pol I transcriptional output via 5-eU metabolic labeling after CX-5461 (10 µM), MTX (1 µM), TPT (50 µM) or CPT (10 µM) treatment for 2 hours. See Figure S2D for details. Statistical significance was calculated using Kruskal-Wallis test followed by Dunn’s multiple comparisons test. \*\*\*\**p*<0.0001, \**p*=0.0220, *ns* = not significant. *n* = 857 (DMSO), 842 (CX-5461), 790 (MTX), 828 (TPT), 213 (CPT) nucleoli. (E) New (4-class) Deep-Phase network outputs for averages of cells treated for 2 hours with varying concentrations of TPT (left) or CPT (right) in activation space. (F) Updated (4-class) Deep-Phase network outputs for averages of cells treated for 2 hours with varying concentrations of MTX in activation space. (G) Dose response curve of nucleolar morphology change and RC_50_ measurements upon 2 hour treatment with varying concentrations of MTX as measured by the new, four class neural network’s vector algebra from the activation space shown in Figure S5F. *n* ≥ 28. (H) Dose response curve of nucleolar morphology change upon 2-hour treatment with varying concentrations of CPT as measured by the updated, four class neural network’s vector algebra from the activation space shown in Figure S5E (right). *n* ≥ 302 cells. (I) Images of HeLa TetR-Cas9 cells, doxycycline-inducible cell lines with control sgRNA (top) or TOP1-targeting sgRNA (bottom) after 6 days of doxycycline treatment (see Figure 5G). Cells were fixed and immunostained with two nucleolar marker antibodies (RPA194, FC and NPM1, GC) and TOP1 antibody to assess knockout efficiency as quantified in Figure 4H. Scale bar = 20 µm. (J) Activation space for a 4-class nucleolar network trained on drug-treated HeLa TetR-Cas9 cells lacking a DFC marker to enable classification of nucleolar morphology based on TOP1 antibody intensity (see Methods). (K) Confusion matrix of the 4-class nucleolar network trained on drug-treated HeLa TetR-Cas9 cells lacking a DFC marker. (L) Classifications of doxycycline (Dox)- or topotecan (TPT)-treated HeLa TetR-Cas9 cells by the four-class nucleolar Deep-Phase network trained on this cell line using all three nucleolar markers that does not allow for sorting based on knockdown efficiency (see Methods). Fractions were calculated from SoftMax function outputs. *n* ≥ 527 cells. (M) Representative confocal microscopy images of HeLa TetR-Cas9 cells with nucleolar marker immunostaining and containing doxycycline-inducible non-targeting (NT) sgRNAs before (left) and after (right) doxycycline treatment (1 µg/mL) for 6 days. Scale bar = 5µm.

**Figure S6.**
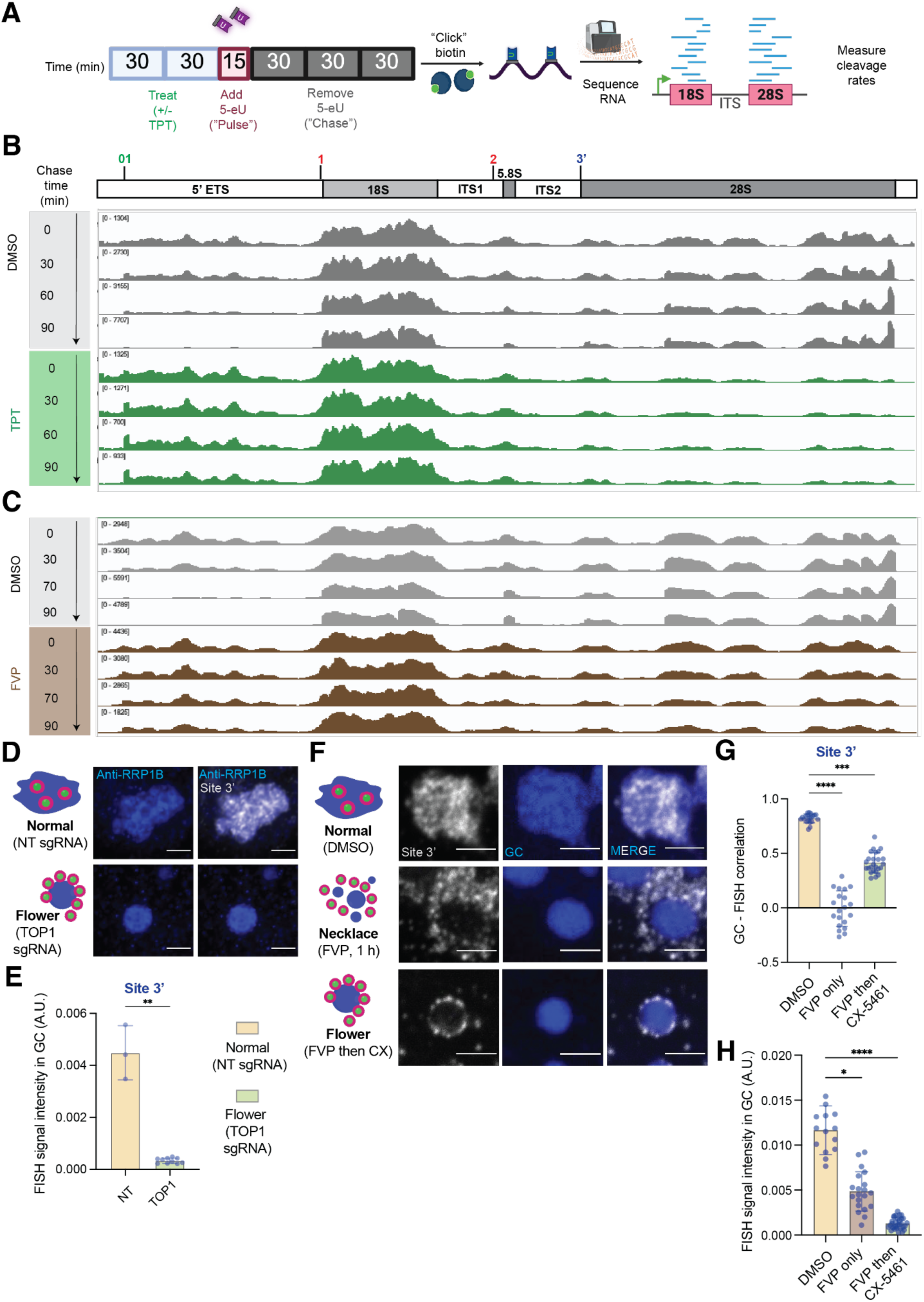
Mechanistic studies of the flower morphology explain how abundance of specific rRNA intermediates shape the nucleolus, related to Figures 6 and 7. (A) Adapted cartoon schematic of the 5-ethynyluridine (5-eU) pulse-chase labeling paradigm to measure pre-rRNA cleavage steps and rates as shown in Figure 6A^94^. (B) Drop in coverage of 5eU-seq reads in the second half of the rRNA transcript in TPT-treated cells. Second of two experimental replicates, see Figure 6F for the first replicate. (C) Previously reported 5eU-seq reads of DMSO-versus FVP-treated cells^94^. (D) Representative confocal microscopy images of RNA FISH experiments targeting site 3’ in HeLa TetR-Cas9 cells, doxycycline-inducible cell lines with control non-targeting sgRNA (top) or TOP1-targeting sgRNA (bottom) after 6 days of doxycycline treatment. Scale bar = 2.5 µm. FISH channel intensity in the bottom image (TOP1 sgRNA) was normalized to the one from the top image (NT sgRNA). (E) Quantification of FISH signal for specific pre-rRNA junctions in HeLa TetR-Cas9 cells after 6 days of doxycycline treatment, see Figures 5G and S6B. Statistical significance was calculated using a Mann-Whitney *U* test. \*\**p*=0.007. (F) Representative min-max confocal microscopy images of RNA FISH experiments targeting site 3’ in the three color nucleolar U-2 OS line. Cells were treated with DMSO (0.05%, 2 hours), FVP (1 µM, 1 hour), or with FVP (1 µM, 1 hour) followed by a wash (2 minutes), and then transcription inhibition with CX-5461 (1 µM, 1 hour). See Figures 7B-C. (G) Quantification of RNA FISH-GC correlation for site 3’ from images shown in Figure S6F (see Methods). Bar height represents the median and error bars represent standard deviation. Statistical significance was calculated using Kruskal-Wallis test followed by Dunn’s multiple comparisons test. \*\*\*\**p*<0.0001, \*\*\**p*=0.0001. *n* = 20 (DMSO and FVP only), 25 (FVP then CX-5461). (H) Quantification of RNA FISH signal intensity in the GC for site 3’ from images shown in Figure S6F (see Methods). Bar height represents the median and error bars represent standard deviation. Statistical significance was calculated using Kruskal-Wallis test followed by Dunn’s multiple comparisons test. \*\*\*\**p*<0.0001, \**p*=0.0346. *n* is the same as in (G).

## SUPPLEMENTAL INFORMATION

Document S1. Tables S1-S8 and supplemental references.

## METHODS

### Cell lines and culture conditions

U-2 OS cells (human female osteosarcoma cells) and HEp-2 (CCL-23) were purchased from ATCC, Lenti-X 293T cells were purchased from Takara, HEK293 cells expressing eYFP-tagged SRRM2 were previously described^121^. All cell lines were grown in a humidified incubator at 37 °C with 5% CO_2_ in Dulbecco’s modified Eagle’s medium (DMEM, Gibco) supplemented with 10% heat-inactivated FBS (Atlanta Biological) and 100 U/mL penicillin-streptomycin (Thermo Fisher). For passaging and imaging, cells were washed with PBS (Gibco) and dissociated from the plate with trypsin-EDTA 0.05% (Fisher Scientific) before being transferred to 96-well glass bottom dishes (Thomas Scientific). For HEK293 cells, dishes were coated with fibronectin prior to plating.

### Plasmid construction

All lentiviral DNA plasmids were generated using the FM5 lentiviral vector containing Ubiquitin C promoter (kind gift from David Sanders). DNA encoding mTagBFP2-NPM1 was PCR-amplified using the Q5 Hot Start High-Fidelity Master Mix (NEB) with primers synthesized by IDT. PCR products were inserted into the linearized vector using the In-Fusion HD cloning kit (Takara) per manufacturer’s instructions. Plasmid inserts were confirmed by Sanger sequencing. RPA16-GFP and Nop56-mCherry plasmids were a kind gift from David Sanders and were previously reported^122^.

### Lentiviral transduction

To create stably expressing U-2 OS lines containing constructs of interest, lentivirus was produced in Lenti-X 293T cells. Specifically, cells were grown to ∼50% confluency in 6-well plates and transfected with the FM5 aforementioned plasmids alongside helper plasmids PSP and VSVG (kind gift from Marc Diamond lab, UT Southwestern) using FuGENE HD transfection reagent (Promega). After three days, supernatant was harvested and used immediately for transduction of U-2 OS cells plated at ∼40% confluency. Three days after transduction, cells were FACS-sorted with a tight window for each fluorescent protein marker and the resulting polyclonal line was passaged for stable maintenance.

### RSV infection and cyclopamine treatment

HEp-2 cells were seeded at 2 × 10⁴ cells per well in 96-well plates and incubated overnight at 37°C. Cells were then infected with 1 × 10⁴ plaque-forming units (PFU) of RSV-GFP (Strain A2, ViraTree, cat #R125) and incubated for 24 hours. Following infection, cells were treated with cyclopamine (CPM; Selleckchem, S1146), solubilized in DMSO at a stock concentration of 10 mM, and applied at final concentrations as indicated in the manuscript. Control wells received 0.25% DMSO. After 1 hour of incubation at 37°C, cells were fixed with 4% paraformaldehyde (PFA) and stained with Hoechst 33342 in DPBS (1:10000 dilution, Thermo Fisher, 62249) prior to imaging.

### Chemical treatments

All cells were plated for at least 12 hours before chemical treatment. For control wells, cells were treated with 0.25 % DMSO. CX-5461, FVP, BMH-21, Oxaliplatin, DRB, THZ1, camptothecin and doxorubicin were purchased from MedChemExpress. Actinomycin D and alpha-amanitin were purchased from Sigma. Pladienolide B was purchased from Tocris Bioscience. Cyclopamine was purchased from ThermoFisher Scientific. All compounds were purchased as powders and resuspended to 10 mM stock concentration, except for Actinomycin D (resuspended to 1 mg/mL), Pladienolide B (resuspended to 1 mM) and alpha-amanitin (1 mg/mL). CX-5461 was resuspended in 50 mM sodium phosphate (pH 3.5), oxaliplatin and alpha-amanitin were resuspended in water, and the remaining compounds were resuspended in DMSO. For the RNA-binding chemical pilot screen, commercial vendors as well as stock resuspension concentrations and media are listed in Table S7. Final chemical treatment concentrations and duration are noted throughout the text and figures.

### Preparation and imaging of cells for neural network processing

Following drug treatment, cells were fixed in 4% paraformaldehyde (diluted in DPBS from a 16% paraformaldehyde stock, Electron Microscopy Science) for 15 minutes at room temperature and washed three times for 5 minutes each in DPBS. Nuclei were stained with the SiR-DNA kit (Spirochrome, CY-SC007) at a 1:5000 dilution in DPBS for at least 3 hours or with Hoechst 33342 (Thermo Fisher Scientific) at a 1:10,000 dilution in DPBS for 30 minutes. Cells were imaged on a Yokogawa CSU-W1 spinning-disc confocal microscope with a Nikon CFI Plan Apo VX 60X water immersion objective (numerical aperture, 1.20) with a Nikon Water Immersion Dispenser (WID) on a Nikon Eclipse Ti2 body using an ORCA-Fusion BT back-thinned high quantum efficiency sCMOS camera. For the three-color nucleolar line, mTagBFP2-NPM1, RPA16-GFP, Nop56-mCherry, and SiR-DNA were imaged using 405, 488, 561 and 647 nm laser lines, respectively. In the HEK293 eYFP-tagged SRRM2 line, 405 and 647 nm lasers were used to image SRRM2 and SiR-DNA, respectively. In RSV-infected HEp-2 cells, 405 and 488 nm laser lines were used to image Hoechst and RSV-GFP, respectively. For TPT-induced morphology validation in wild-type U-2 OS and wild-type HeLa cells, Hoechst, anti-RPA194, anti-Fibrillarin, and anti-NPM1 were imaged using 405, 488, 561 and 647 nm laser lines. For dose-response validation in wild-type U-2 OS and for TOP1 KO in CRISPR HeLa cell lines, anti-RRP1B, anti-RPA194, anti-Fibrillarin, and SiR-DNA were imaged using 405, 488, 561 and 647 nm laser lines. For the updated Deep-Phase network in CRISPR HeLa cell lines that enabled binning by TOP1 antibody intensity, the anti-Fibrillarin stain was replaced with anti-TOP1. Using Nikon’s JOBS software module, an automated image acquisition protocol was developed where four to nine 3×3 images per well taken (608 x 608 µm each), depending on cell type and confluence. Images were batch-denoised (4 channels) using Nikon Elements Denoise.ai module and batch-deconvolved (20 iterations) with the Richardson-Lucy algorithm.

### Deep-Phase architecture, training, and usage

Deep-Phase is adapted from a pre-trained ResNet architecture^67^ by replacing the last fully connected layer with two fully connected, linear layers. The additional hidden layer has two neurons, which are used to monitor and display the response of each cell in activation space. The number of outputs of the last layer is chosen to match the number of cell phenotypes required. While restricting the latent space to two dimensions and omitting a non-linear activation function likely decreases the power of the overall network, linearity is imperative to generate a response metric which scales directly with phenotypic changes. Furthermore, the network is sufficient to discriminate between the few classes used here as demonstrated by the high accuracy.

The trained ResNet expects input images to be 224 by 224 pixel RGB images, normalized between 0 and 1 prior to scaling to a mean of ∼0.45 and standard deviation of ∼0.225. Initially, we experimented with freezing training on all but the last fully connected layers and therefore we provided input images adhering to these constraints. For each image, only the filename and cell locations are needed. To adapt Deep-Phase to different experiments, users can adjust how cell finding is performed and which channels to map into the RGB channels. We note that the near-uniform cellular responses to each small molecule treatment allowed us to use treatment conditions as a surrogate label for single-cell morphology, facilitating the rapid generation of large, automatically labeled training datasets. This approach minimized labeling bias and enhanced the performance and generalizability of the convolutional neural networks.

To train and evaluate a Deep-Phase network, we begin with parsing individual cell locations and filenames from the output table from a CellProfiler pipeline^123^ in our purpose-built python software. For each cell, a 384 by 384 pixel area is cropped from the input image with the x,y position at its center. The image is resized to a 256 by 256 pixel image and the fluorescence channels are used to create an RGB composite. First, each channel is normalized between the minimum and maximum intensity within each cropped region. The user specifies how each channel is mapped to the red, green, and blue channels. The 256 by 256 RGB images are split into a testing and training set. The test set is utilized in evaluation mode, where the center 224 pixels are cropped and the image is normalized for input to the ResNet. The training set is augmented further in the python software with a 224 crop at a random position, horizontal and vertical flips, and, if desired, rotations and gaussian noise. The random rotation and gaussian noise were optimized as hyperparameters.

We experimented with several hyperparameters and, based on lowest test set loss, arrived at the following values for training: a ResNet50 initialized to imagenet1k_v2 weights, with all layers left unfrozen, and random rotations limited to 10 degrees and gaussian noise of 0.001, as they were found to optimize robustness and training performance. When running in inference mode, a forward hook is registered on the final hidden layer to monitor the activations on each neuron. Additionally, the logit output of the network is softmax-transformed into a probability estimate for each class. The activations and class probabilities are appended to the CellProfiler datatable for further processing and analysis. To quantify response of a specific cell, the mean activation of untreated (DMSO) and treated (drug) cells used in training are utilized as standards. The response of a cell is then the relative, projected distance between untreated and treated standards in activation space. Importantly, the treated condition is chosen to match each perturbation, e.g. CX-5461 for cap forming compounds and FVP for necklace forming compounds.

As mentioned above, the network can be tailored to specific experiments by adjusting the cell finding algorithm and mapping of color channels. For the original 3-class nucleolus network, cell center positions were found using CellProfiler v4.2.6. Briefly, the SiR-DNA channel was selected and primary cell objects were found using global, minimum cross-entropy thresholding, retaining objects with diameters between 120 and 240 pixels. Putative cells were further filtered to have a circularity between 0.7 and 1 to avoid cells undergoing division. Mapping the 4 fluorescence channels to RGB was performed by assigning the FC, DFC, and GC channels to red, green and blue channels, respectively, with intensity scaled by 0.75. The SiR-DNA channel is added to each color channel with a scaling of 0.25. This balance was chosen to emphasize the nucleolar components without completely discarding the nuclear images. The updated, 4-class nucleolus network that included TPT-treated cells was set up identically except the addition of a flower phenotype for a total of four possible classes.

When applied to CRISPR HeLa cells with the goal of correlating TOP1 abundance to nucleolar morphology, because of the heterogeneity in TOP1 knockout efficiency, the network was tailored to account for the loss of the DFC image as a channel was required for TOP1 antibody staining. While the fluorescence channels can be adjusted between training and inference, we find that performance generally degrades compared to a network trained on the same components. As such, SiR-DNA, FC and GC channels were assigned to red, green and blue for input to the network. Cell locations were found as above.

The nuclear speckle network used the same cell finding algorithm, but the only inputs to the network were the SiR-DNA and eYFP-SRRM2 channels, which were assigned to red and green respectively; the blue channel was set to 0.

Finally, the cytoplasmic RSV condensate images required additional processing prior to applying Deep-Phase. Cell cytoplasm was found using the cyto2 model of Cellpose v2.3.2^124,125^ with the RSV-GFP channel and Hoecsht as the nucleus channel with an expected diameter of 300 pixels. Cellpose was run again with the nuclei model on the Hoechst image and an object diameter of 100 pixels. The nuclear regions were subtracted from the cells to obtain a mask of only the cytoplasm which was used to mask the RSV-GFP image and determine cell positions. The putative cells were further refined to exclude cells with multiple nuclei and those with cytoplasmic areas of less than 5000 pixels, which occasionally occurred when the cyto2 model found cells without appreciable RSV-GFP signal. For Deep-Phase training and inference, the nuclei-masked RSV-GFP image was assigned to all color channels.

### Metabolic labeling and imaging of nascent RNA with 5-eU

Three-color nucleolar U-2 OS cells were treated with indicated drug concentrations or DMSO for 1 hour and 30 minutes at 37°C and 5% CO_2_ prior to addition of 1 mM of 5-ethynyl uridine (5-eU, Cedarlane, CLK-N002-10). After a 15 minute incubation with 5-eU and drugs (“pulse”), cells were washed twice with DPBS and treated with drugs only for an additional 15 minutes (“chase”). Drug-containing media was then removed and cells were fixed in 4% paraformaldehyde (diluted in DPBS from a 16% paraformaldehyde stock, Electron Microscopy Science) for 15 minutes at room temperature. After removing the fixative, cells were washed twice in DPBS and permeabilized in 0.5% Triton™ X-100 (Thermo Fisher) in DPBS for 15 minutes at room temperature. Cells were again washed twice in DPBS and treated with blocking solution containing 2% RNase-free bovine serum albumin (BSA, Avantor, 97061) for 1 hour before the addition of RPA194 Antibody (C-1) Alexa Fluor® 488 (sc-48385) diluted 1:400 in blocking solution. Cells were incubated for 1 hour at room temperature. Next, 5-eU was labeled with the Click-iT™ Plus Alexa Fluor™ 647 Picolyl Azide Toolkit (Thermo Fisher, C10643) per manufacturer’s instructions. Cells were imaged on a Yokogawa CSU-W1 spinning-disc confocal microscope with a Nikon 60X oil-immersion Plan Apo λD objective (numerical aperture, 1.42) on a Nikon Eclipse Ti2 body using an ORCA-Fusion BT back-thinned high quantum efficiency sCMOS camera. mTagBFP2-NPM1, anti-RPA194, Nop56-mCherry, and 5-eU were imaged using 405, 488, 561 and 647 nm laser lines, respectively.

### Immunofluorescence and imaging

Cells were fixed in 4% paraformaldehyde (diluted in DPBS from a 16% paraformaldehyde stock, Electron Microscopy Science) for 15 minutes at room temperature and then washed three times in DPBS for 5 minutes each. Cells were incubated in 5% normal goat serum (Vector Laboratories) and 0.3% Triton™ X-100 (Thermo Fisher) in DPBS for 1 hour at room temperature. Primary antibodies were diluted in 0.3% Triton™ X-100 and 1% BSA (Avantor, 97061) in DPBS and incubated overnight at room temperature. Phospho-Rpb1 CTD Ser2/5 antibody (rabbit, Cell Signaling Technology, 4735) was used at 1:200 dilution, TOP1 antibody (rabbit, Thermo Fisher, MA5-32228) was used at 1:200 dilution, and nucleolin antibody (mouse, Thermo Fisher, 39-6400) was used at 1:100 dilution. Cells were washed three times in DPBS for 5 minutes each prior to secondary antibody incubation with goat anti-rabbit or goat anti-mouse Alexa Fluor™ 647 (Thermo Fisher, A-21245 and A-21235, respectively). HEK293 eYFP-tagged SRRM2 line was additionally stained with Hoechst 33342 in DPBS (1:10000 dilution, Thermo Fisher, 62249). The three-color nucleolar U-2 OS line was imaged on a Yokogawa CSU-W1 spinning-disc confocal microscope with a Nikon 60X oil-immersion Plan Apo λD objective (numerical aperture, 1.42) on a Nikon Eclipse Ti2 body using an ORCA-Fusion BT back-thinned high quantum efficiency sCMOS camera. mTagBFP2-NPM1, RPA194, Nop56-mCherry, and antibodies were imaged using 405, 488, 561 and 647 nm laser lines, respectively. The HEK293 eYFP-tagged SRRM2 line was imaged on a Yokogawa CSU-W1 spinning-disc confocal microscope with a Nikon CFI Plan Apo VX 60X water immersion objective (numerical aperture, 1.20) with a Nikon Water Immersion Dispenser (WID) on a Nikon Eclipse Ti2 body using an ORCA-Fusion BT back-thinned high quantum efficiency sCMOS camera. Hoechst 33342, eYFP and Phospho-Rpb1 CTD Ser2/5 antibody were imaged with a 405, 488, and 647 nm laser line, respectively.

### RNA-binding chemical pilot screen

Cells were plated in a 96-well plate at least 12 hours before treatment. Each 96 well plate contained duplicate wells treated with DMSO (0.25%), CX-5461 (10 µM) and FVP (6.25 µM) as well as the 23 purchased compounds (Table S7) at 1, 10, and 50 µM concentrations in duplicate. Cells were treated for 2 hours prior to being subjected to fixation, nuclear staining, and imaging for Deep-Phase analysis as described above.

### 5eU-seq with topotecan and data analysis

The experiment was conducted as previously described^94^. Briefly, the three-color nucleolar U-2 OS line was seeded in a 6 well plate, with one well used per condition. Each condition had two biological replicates. Cells treated with 50 µM of topotecan or DMSO in DMEM for 1 hour prior to addition of 1 mM of 5-ethynyl uridine (5-eU, Cedarlane, CLK-N002-10). After a 15 minute incubation with 5-eU and topotecan or DMSO (“pulse”), cells were washed twice with DPBS containing 10 mM uridine (Sigma, U6381) and lysed immediately (for 0 min chase) or treated with 50 µM of topotecan or DMSO in DMEM containing 10 mM uridine for an additional 30, 60, or 90 minutes prior to lysing. Cells were lysed in buffer RLT (Qiagen, 79216). Total RNA was isolated and DNAse-treated prior to biotinylation and streptavidin-bead capture of 5eU-labelled RNA as previously described. RNA-seq libraries were generated as previously described^94^ and paired-end sequencing (75×75) was performed on an Illumina MiSeq sequencer.

A custom snakemake pipeline was used to perform analysis of the 5eU-seq reads (https://github.com/SoftLivingMatter/5eU-seq-pipelines). Briefly, reads were trimmed, aligned to the rDNA genome (GenBank: U13369.1), and then visualized in Integrative Genome Viewer (IGV)^126^. The fraction cleaved metric was determined by calculating the fraction of sequencing reads spanning (not cleaved) vs. not spanning (cleaved) multiple known rRNA cleavage junctions as previously described^94^. The 18S-to-28S ratio was calculated using bedtools intersect^127^ by calculating the number of reads in either the 18S (3657-5527 nt) vs 28S (7935-12969 nt) regions, respectively, and then plotted using GraphPad Prism v10.1.1.

### Generation of Dox-inducible CRISPR lines

TOP1 and non-targeting (NT) sgRNAs were cloned into CropSeq-purov2 (Addgene 127458) using golden gate cloning and sequence-verified. sgRNA oligonucleotide sequences can be found in Table S8.

For generation of lentivirus, 300k HE293T cells were seeded in 6-well plates. The following day, lentivirus was prepared by transfecting HEK293T cells with 400ng VSV-G envelope plasmid, 1200ng PSP packaging plasmid, and 400ng of an individual sgRNA plasmid using lipofectamine 3000 and following manufacturer’s instructions. Two days following transfection, lentivirus was harvested and filtered.

HeLa TetR-Cas9 cells^128^ (gift from Iain Cheeseman, Whitehead Institute/MIT) grown in Tet-free Fetal Bovine Serum (FBS; Omega Scientific FB-15) were infected with either NT or TOP1 sgRNA lentivirus at an approximate multiplicity of infection of 2. 2 days after virus addition, cells were washed with PBS and media was replaced with media containing 2ug/mL puromycin (Sigma P4512-1MLX10). 3 days later, puromycin-containing media was replaced with media without puromycin for further growth. Complete selection was confirmed by comparison in parallel with non-infected HeLa TetR-Cas9 cells in the presence of puromycin.

To induce Cas9 in HeLa TetR-Cas9 cells with TOP1 sgRNA or NT sgRNA, cells were incubated in the presence of 1ug/mL doxycycline (Clontech NC0424034) for 6 days.

### RNA SABER FISH experiments

The experiments were conducted as described previously^94,129,130^ and probe sequences for rRNA site 3’ were reported previously^94^. All probes were concatemerized with hairpin 25, which was paired with 647 fluorescent oligonucleotides for imaging. Briefly, after drug treatment, cells were fixed in 4% PFA for 10 minutes and washed in PBST for 15 minutes at room temperature. Next, cells were incubated in Whyb solution (2X SCC, 1% Tween-20, and 40% deionized formamide) for 1 hour at 44 °C at 300 rpm, and then left at 43°C for 16 hours at 300 rpm in hybridization solution (2X SCC, 1% Tween-20, 40% deionized formamide, 10% wt/vol dextran sulfate) with 1 µg of concatemerized probe. Following this incubation, cells were washed twice for 30 minutes in Whyb solution at 42 °C and 300 rpm, twice for 5 minutes in 2X SCC and 0.1% Tween-20 at 42 °C and 300 rpm, and finally twice for 1 minute in PBST. Cells were then incubated in 0.4 µM fluorescent imager probes in PBST at 37 °C for 10 minutes. After three washes at 37°C in PBST for 5 minutes and one wash at room temperature in PBS, cells were blocked in 2% BSA and stained with anti-RRP1B as described previously. Blocking and staining steps contained murine RNase inhibitor (1:200, New England Biolabs, M0314). Finally, cell nuclei were stained with YOYO-1 iodide for 1 hour (Thermo Fisher Scientific, Y3601).

### Quantitative image analysis for 5-eU metabolic labeling, immunofluorescence, TOP1 partitioning, and SABER-FISH datasets

All analyses were done in CellProfiler v4.2.6^123^ and all quantified data was plotted in GraphPad Prism v10.1.1.

For quantifying 5-eU metabolic labeling signal as a readout of Pol I transcriptional output, individual GCs were first identified using the IdentifyPrimaryObjects module and the mTagBFP2-NPM1 channel with two-class global Otsu thresholding. Then, the Nop56-mCherry channel was first enhanced using the EnhanceOrSuppressFeatures module with Speckles as the feature type and 40 as the feature size. This enhanced image was used as input to identify individual DFCs using the IdentifyPrimaryObjects module with three-class global Otsu thresholding. Pixels in the middle intensity class were assigned to the background. 5-eU intensity was measured by using the MeasureObjectIntensity module and selecting the 5-eU channel to be measured in the previously identified DFCs. Finally, individual DFCs were assigned to their corresponding GCs by using the RelateObjects module, with GCs as the parent and DFCs as the child objects.

For quantifying anti-phospho-Rpb1 CTD Ser2/5 intensity as a proxy for active Pol II, or anti-TOP1 to assess TOP1 knockout efficiency, individual nuclei were first identified using the IdentifyPrimaryObjects module and the Hoechst channel with three-class global Otsu thresholding. Pixels from the middle intensity class were assigned to the foreground. Then, antibody intensity was measured by using the MeasureObjectIntensity module and selecting the antibody channel to be measured in the previously identified nuclei.

For quantifying nucleolar-nucleoplasmic partitioning of anti-TOP1, individual nuclei were first identified using the IdentifyPrimaryObjects module and the TOP1 channel with two-class global Otsu thresholding. Then, individual GCs were identified using the IdentifyPrimaryObjects module and the same thresholding settings. The nucleoplasm was identified by using the IdentifyTertiaryObjects module with nuclei as the larger and GCs as the smaller identified objects. TOP1 intensity was measured using the MeasureObjectIntensity module in the nucleoplasm and GCs. Finally, individual GCs were assigned to their corresponding nucleoplasm by using the RelateObjects module, with nucleoplasm as the parent and GCs as the child objects. Partitioning (K) was calculated by dividing the average TOP1 intensity in GCs per cell by the TOP1 intensity in the nucleoplasm of the corresponding nucleus.

For measuring GC correlation and GC intensity of site 3’ pre-rRNA from SABER-FISH experiments, we adapted a previously reported pipeline^94^. Specifically, individual nuclei locations were found with IdentifyPrimaryObjects using a global Otsu threshold on the YOYO-1 stained image after Gaussian smoothing. The nuclear objects were used to mask the GC image to detect nucleoli using an adaptive Otsu threshold for objects between 15 and 300 pixels without dividing clumped objects. The nucleoli were then dilated by 20 pixels to form the basis for determining colocalization later. The FISH and GC images, corresponding to the FISH fluorescent imager probe and mTagBFP2-NPM1 or anti-RRP1B, respectively, were background-subtracted using the 5th quantile intensity of each channel. The colocalization of the background subtracted images, size and intensity were measured for each nucleolus, grouped into a given nucleus. Using the dilated region around nucleoli can increase sensitivity of correlation measurements by reducing the amount of background from the nuclear space.

### Statistical analysis

All statistical analyses were conducted in GraphPad Prism v10.1.1 software (GraphPad). Statistical significance for functional assays, including 5-eU and phosphorylated PolII antibody intensity as well as for TOP1 antibody staining to assess knockout efficiency and for RNA FISH experiments with drug-treated cells was calculated using a Kruskal-Wallis test followed by Dunn’s multiple comparisons test. Statistical significance for TOP1 partitioning change during drug treatment was calculated using two-tailed Mann-Whitney *U* tests. Statistical significance for RNA FISH in TOP1 knockout cells was conducted using a Mann-Whitney *U* test. Corresponding *p* values and size of *n* are noted in figure legends.

## REFERENCES

1. Banani, S.F., Lee, H.O., Hyman, A.A., and Rosen, M.K. (2017). Biomolecular condensates: organizers of cellular biochemistry. Nat Rev Mol Cell Biol 18, 285–298.

2. Shin, Y., and Brangwynne, C.P. (2017). Liquid phase condensation in cell physiology and disease. Science 357. 10.1126/science.aaf4382.

3. Holehouse, A.S., and Pappu, R.V. (2018). Functional Implications of Intracellular Phase Transitions. Biochemistry 57, 2415–2423.

4. Alberti, S., Gladfelter, A., and Mittag, T. (2019). Considerations and Challenges in Studying Liquid-Liquid Phase Separation and Biomolecular Condensates. Cell 176, 419–434.

5. Mitrea, D.M., and Kriwacki, R.W. (2016). Phase separation in biology; functional organization of a higher order. Cell Commun Signal 14, 1.

6. Dignon, G.L., Best, R.B., and Mittal, J. (2020). Biomolecular Phase Separation: From Molecular Driving Forces to Macroscopic Properties. Annu Rev Phys Chem 71, 53–75.

7. Schuster, B.S., Regy, R.M., Dolan, E.M., Kanchi Ranganath, A., Jovic, N., Khare, S.D., Shi, Z., and Mittal, J. (2021). Biomolecular Condensates: Sequence Determinants of Phase Separation, Microstructural Organization, Enzymatic Activity, and Material Properties. J Phys Chem B 125, 3441–3451.

8. Holehouse, A.S., and Alberti, S. (2025). Molecular determinants of condensate composition. Mol Cell 85, 290–308.

9. Brangwynne, C.P., Eckmann, C.R., Courson, D.S., Rybarska, A., Hoege, C., Gharakhani, J., Jülicher, F., and Hyman, A.A. (2009). Germline P granules are liquid droplets that localize by controlled dissolution/condensation. Science 324, 1729–1732.

10. Feric, M., Vaidya, N., Harmon, T.S., Mitrea, D.M., Zhu, L., Richardson, T.M., Kriwacki, R.W., Pappu, R.V., and Brangwynne, C.P. (2016). Coexisting Liquid Phases Underlie Nucleolar Subcompartments. Cell 165, 1686–1697.

11. Maharana, S., Wang, J., Papadopoulos, D.K., Richter, D., Pozniakovsky, A., Poser, I., Bickle, M., Rizk, S., Guillén-Boixet, J., Franzmann, T.M., et al. (2018). RNA buffers the phase separation behavior of prion-like RNA binding proteins. Science 360, 918–921.

12. Strom, A.R., Emelyanov, A.V., Mir, M., Fyodorov, D.V., Darzacq, X., and Karpen, G.H. (2017). Phase separation drives heterochromatin domain formation. Nature 547, 241–245.

13. Sabari, B.R., Dall’Agnese, A., Boija, A., Klein, I.A., Coffey, E.L., Shrinivas, K., Abraham, B.J., Hannett, N.M., Zamudio, A.V., Manteiga, J.C., et al. (2018). Coactivator condensation at super-enhancers links phase separation and gene control. Science 361. 10.1126/science.aar3958.

14. Boeynaems, S., Holehouse, A.S., Weinhardt, V., Kovacs, D., Van Lindt, J., Larabell, C., Van Den Bosch, L., Das, R., Tompa, P.S., Pappu, R.V., et al. (2019). Spontaneous driving forces give rise to protein-RNA condensates with coexisting phases and complex material properties. Proc Natl Acad Sci U S A 116, 7889–7898.

15. Li, P., Banjade, S., Cheng, H.-C., Kim, S., Chen, B., Guo, L., Llaguno, M., Hollingsworth, J.V., King, D.S., Banani, S.F., et al. (2012). Phase transitions in the assembly of multivalent signalling proteins. Nature 483, 336–340.

16. Nott, T.J., Petsalaki, E., Farber, P., Jervis, D., Fussner, E., Plochowietz, A., Craggs, T.D., Bazett-Jones, D.P., Pawson, T., Forman-Kay, J.D., et al. (2015). Phase transition of a disordered nuage protein generates environmentally responsive membraneless organelles. Mol Cell 57, 936–947.

17. Elbaum-Garfinkle, S., Kim, Y., Szczepaniak, K., Chen, C.C.-H., Eckmann, C.R., Myong, S., and Brangwynne, C.P. (2015). The disordered P granule protein LAF-1 drives phase separation into droplets with tunable viscosity and dynamics. Proc Natl Acad Sci U S A 112, 7189–7194.

18. Bracha, D., Walls, M.T., Wei, M.-T., Zhu, L., Kurian, M., Avalos, J.L., Toettcher, J.E., and Brangwynne, C.P. (2019). Mapping Local and Global Liquid Phase Behavior in Living Cells Using Photo-Oligomerizable Seeds. Cell 176, 407.

19. Riback, J.A., Zhu, L., Ferrolino, M.C., Tolbert, M., Mitrea, D.M., Sanders, D.W., Wei, M.-T., Kriwacki, R.W., and Brangwynne, C.P. (2020). Composition-dependent thermodynamics of intracellular phase separation. Nature 581, 209–214.

20. Shimobayashi, S.F., Ronceray, P., Sanders, D.W., Haataja, M.P., and Brangwynne, C.P. (2021). Nucleation landscape of biomolecular condensates. Nature 599, 503–506.

21. Rai, A.K., Chen, J.-X., Selbach, M., and Pelkmans, L. (2018). Kinase-controlled phase transition of membraneless organelles in mitosis. Nature 559, 211–216.

22. Banani, S.F., Rice, A.M., Peeples, W.B., Lin, Y., Jain, S., Parker, R., and Rosen, M.K. (2016). Compositional Control of Phase-Separated Cellular Bodies. Cell 166, 651–663.

23. Klein, I.A., Boija, A., Afeyan, L.K., Hawken, S.W., Fan, M., Dall’Agnese, A., Oksuz, O., Henninger, J.E., Shrinivas, K., Sabari, B.R., et al. (2020). Partitioning of cancer therapeutics in nuclear condensates. Science 368, 1386–1392.

24. Banani, S.F., Afeyan, L.K., Hawken, S.W., Henninger, J.E., Dall’Agnese, A., Clark, V.E., Platt, J.M., Oksuz, O., Hannett, N.M., Sagi, I., et al. (2022). Genetic variation associated with condensate dysregulation in disease. Dev Cell 57, 1776–1788.e8.

25. Li, Y.R., King, O.D., Shorter, J., and Gitler, A.D. (2013). Stress granules as crucibles of ALS pathogenesis. J Cell Biol 201, 361–372.

26. Alberti, S., and Dormann, D. (2019). Liquid-Liquid Phase Separation in Disease. Annu Rev Genet 53, 171–194.

27. Visser, B.S., Lipiński, W.P., and Spruijt, E. (2024). The role of biomolecular condensates in protein aggregation. Nat Rev Chem 8, 686–700.

28. Jiang, L., and Kang, Y. (2025). Biomolecular condensates: A new lens on cancer biology. Biochim Biophys Acta Rev Cancer 1880, 189245.

29. Boija, A., Klein, I.A., and Young, R.A. (2021). Biomolecular Condensates and Cancer. Cancer Cell 39, 174–192.

30. Mitrea, D.M., Mittasch, M., Gomes, B.F., Klein, I.A., and Murcko, M.A. (2022). Modulating biomolecular condensates: a novel approach to drug discovery. Nat. Rev. Drug Discov. 21, 841–862.

31. Conti, B.A., and Oppikofer, M. (2022). Biomolecular condensates: new opportunities for drug discovery and RNA therapeutics. Trends Pharmacol Sci 43, 820–837.

32. Eftekharzadeh, B., Mayfield, A., Kauffman, M.G., and Reilly, J.F. (2024). Drug Discovery for Diseases with High Unmet Need Through Perturbation of Biomolecular Condensates. J Mol Biol 436, 168855.

33. Mullard, A. (2019). Biomolecular condensates pique drug discovery curiosity. Nat Rev Drug Discov. 10.1038/d41573-019-00069-w.

34. Fang, M.Y., Markmiller, S., Vu, A.Q., Javaherian, A., Dowdle, W.E., Jolivet, P., Bushway, P.J., Castello, N.A., Baral, A., Chan, M.Y., et al. (2019). Small-Molecule Modulation of TDP-43 Recruitment to Stress Granules Prevents Persistent TDP-43 Accumulation in ALS/FTD. Neuron 103, 802–819.e11.

35. Boyd, J.D., Lee, P., Feiler, M.S., Zauur, N., Liu, M., Concannon, J., Ebata, A., Wolozin, B., and Glicksman, M.A. (2014). A high-content screen identifies novel compounds that inhibit stress-induced TDP-43 cellular aggregation and associated cytotoxicity. J Biomol Screen 19, 44–56.

36. Chandrasekaran, S.N., Ceulemans, H., Boyd, J.D., and Carpenter, A.E. (2021). Image-based profiling for drug discovery: due for a machine-learning upgrade? Nat. Rev. Drug Discov. 20, 145–159.

37. Danuser, G. (2011). Computer vision in cell biology. Cell 147, 973–978.

38. Vasan, R., Ferrante, A.J., Borensztejn, A., Frick, C.L., Garrison, P., Gaudreault, N., Mogre, S.S., Mohammed, F.S., Morris, B., Pires, G.G., et al. (2025). Interpretable representation learning for 3D multi-piece intracellular structures using point clouds. Nat Methods 22, 1531–1544.

39. Shen, D., Zhu, Q., Pang, X., Yang, X., Pan, D., Zhang, M., Li, Y., Sun, Z., Fang, L., Chen, W., et al. (2025). PB-scope: Contrastive learning of dynamic processing body formation reveals undefined mechanisms of approved compounds. bioRxiv. 10.1101/2025.06.14.659731.

40. Gut, G., Herrmann, M.D., and Pelkmans, L. (2018). Multiplexed protein maps link subcellular organization to cellular states. Science 361. 10.1126/science.aar7042.

41. Spitzer, H., Berry, S., Donoghoe, M., Pelkmans, L., and Theis, F.J. (2023). Learning consistent subcellular landmarks to quantify changes in multiplexed protein maps. Nat. Methods 20, 1058–1069.

42. Krentzel, D., Shorte, S.L., and Zimmer, C. (2023). Deep learning in image-based phenotypic drug discovery. Trends Cell Biol. 33, 538–554.

43. LeCun, Y., Bengio, Y., and Hinton, G. (2015). Deep learning. Nature 521, 436–444.

44. Moen, E., Bannon, D., Kudo, T., Graf, W., Covert, M., and Van Valen, D. (2019). Deep learning for cellular image analysis. Nat Methods 16, 1233–1246.

45. Thiry, M., and Lafontaine, D.L.J. (2005). Birth of a nucleolus: the evolution of nucleolar compartments. Trends Cell Biol 15, 194–199.

46. Pederson, T. (2011). The nucleolus. Cold Spring Harb Perspect Biol 3. 10.1101/cshperspect.a000638.

47. Shav-Tal, Y., Blechman, J., Darzacq, X., Montagna, C., Dye, B.T., Patton, J.G., Singer, R.H., and Zipori, D. (2005). Dynamic sorting of nuclear components into distinct nucleolar caps during transcriptional inhibition. Mol Biol Cell 16, 2395–2413.

48. Burger, K., Mühl, B., Harasim, T., Rohrmoser, M., Malamoussi, A., Orban, M., Kellner, M., Gruber-Eber, A., Kremmer, E., Hölzel, M., et al. (2010). Chemotherapeutic drugs inhibit ribosome biogenesis at various levels. J. Biol. Chem. 285, 12416–12425.

49. Schmidt, H.B., Jaafar, Z.A., Wulff, B.E., Rodencal, J.J., Hong, K., Aziz-Zanjani, M.O., Jackson, P.K., Leonetti, M.D., Dixon, S.J., Rohatgi, R., et al. (2022). Oxaliplatin disrupts nucleolar function through biophysical disintegration. Cell Rep 41, 111629.

50. Alley, K.R., Wyatt, K.M., Fries, A.C., and DeRose, V.J. (2025). Expansion Microscopy Provides Nanoscale Insight into Nucleolar Reorganization and Nuclear Foci Formation during Nucleolar Stress. ACS Chem Biol 20, 1232–1246.

51. Nicolas, E., Parisot, P., Pinto-Monteiro, C., de Walque, R., De Vleeschouwer, C., and Lafontaine, D.L.J. (2016). Involvement of human ribosomal proteins in nucleolar structure and p53-dependent nucleolar stress. Nat. Commun. 7, 11390.

52. Stamatopoulou, V., Parisot, P., De Vleeschouwer, C., and Lafontaine, D.L.J. (2018). Use of the iNo score to discriminate normal from altered nucleolar morphology, with applications in basic cell biology and potential in human disease diagnostics. Nat. Protoc. 13, 2387–2406.

53. Farley-Barnes, K.I., McCann, K.L., Ogawa, L.M., Merkel, J., Surovtseva, Y.V., and Baserga, S.J. (2018). Diverse Regulators of Human Ribosome Biogenesis Discovered by Changes in Nucleolar Number. Cell Rep. 22, 1923–1934.

54. Ogawa, L.M., Buhagiar, A.F., Abriola, L., Leland, B.A., Surovtseva, Y.V., and Baserga, S.J. (2021). Increased numbers of nucleoli in a genome-wide RNAi screen reveal proteins that link the cell cycle to RNA polymerase I transcription. Mol. Biol. Cell 32, 956–973.

55. Potapova, T.A., Unruh, J.R., Conkright-Fincham, J., Banks, C.A.S., Florens, L., Schneider, D.A., and Gerton, J.L. (2023). Distinct states of nucleolar stress induced by anticancer drugs. Elife 12. 10.7554/eLife.88799.

56. Sheu-Gruttadauria, J., Yan, X., Stuurman, N., Vale, R.D., and Floor, S.N. (2024). Nucleolar dynamics are determined by the ordered assembly of the ribosome. bioRxiv. 10.1101/2023.09.26.559432.

57. Lafontaine, D.L., and Tollervey, D. (2000). Synthesis and assembly of the box C+D small nucleolar RNPs. Mol. Cell. Biol. 20, 2650–2659.

58. Okuwaki, M., Saito, S., Hirawake-Mogi, H., and Nagata, K. (2021). The interaction between nucleophosmin/NPM1 and the large ribosomal subunit precursors contribute to maintaining the nucleolar structure. Biochim. Biophys. Acta Mol. Cell Res. 1868, 118879.

59. Haddach, M., Schwaebe, M.K., Michaux, J., Nagasawa, J., O’Brien, S.E., Whitten, J.P., Pierre, F., Kerdoncuff, P., Darjania, L., Stansfield, R., et al. (2012). Discovery of CX-5461, the First Direct and Selective Inhibitor of RNA Polymerase I, for Cancer Therapeutics. ACS Med. Chem. Lett. 3, 602–606.

60. Drygin, D., Lin, A., Bliesath, J., Ho, C.B., O’Brien, S.E., Proffitt, C., Omori, M., Haddach, M., Schwaebe, M.K., Siddiqui-Jain, A., et al. (2011). Targeting RNA polymerase I with an oral small molecule CX-5461 inhibits ribosomal RNA synthesis and solid tumor growth. Cancer Res. 71, 1418–1430.

61. Reynolds, R.C., Montgomery, P.O., and Hughes, B. (1964). NUCLEOLAR “CAPS” PRODUCED BY ACTINOMYCIN D. Cancer Res 24, 1269–1277.

62. Schoefl, G.I. (1964). THE EFFECT OF ACTINOMYCIN D ON THE FINE STRUCTURE OF THE NUCLEOLUS. J Ultrastruct Res 10, 224–243.

63. Sedlacek, H.H. (2001). Mechanisms of action of flavopiridol. Crit. Rev. Oncol. Hematol. 38, 139–170.

64. Caudron-Herger, M., Pankert, T., Seiler, J., Németh, A., Voit, R., Grummt, I., and Rippe, K. (2015). Alu element-containing RNAs maintain nucleolar structure and function. EMBO J. 34, 2758–2774.

65. Abraham, K.J., Khosraviani, N., Chan, J.N.Y., Gorthi, A., Samman, A., Zhao, D.Y., Wang, M., Bokros, M., Vidya, E., Ostrowski, L.A., et al. (2020). Nucleolar RNA polymerase II drives ribosome biogenesis. Nature 585, 298–302.

66. Serafim, R.B., Volegova, M., Harlow, M., Banerjee, U., Cardoso, C., Das, S., Sharma, B., Zentner, G.E., Scacheri, P.C., Zhang, T., et al. (2025). CDK12/13 inhibition disrupts nucleolar morphology and promotes aberrant expression of IGS transcripts. bioRxiv. 10.1101/2025.03.01.640972.

67. He, K., Zhang, X., Ren, S., and Sun, J. (2015). Deep residual learning for image recognition. arXiv [cs.CV]. 10.48550/ARXIV.1512.03385.

68. Chen, S., Ding, C., Liu, M., Cheng, J., and Tao, D. (2023). CPP-Net: Context-Aware Polygon Proposal Network for Nucleus Segmentation. IEEE Trans Image Process 32, 980– 994.

69. Greenwald, N.F., Miller, G., Moen, E., Kong, A., Kagel, A., Dougherty, T., Fullaway, C.C., McIntosh, B.J., Leow, K.X., Schwartz, M.S., et al. (2022). Whole-cell segmentation of tissue images with human-level performance using large-scale data annotation and deep learning. Nat Biotechnol 40, 555–565.

70. Peltonen, K., Colis, L., Liu, H., Trivedi, R., Moubarek, M.S., Moore, H.M., Bai, B., Rudek, M.A., Bieberich, C.J., and Laiho, M. (2014). A targeting modality for destruction of RNA polymerase I that possesses anticancer activity. Cancer Cell 25, 77–90.

71. Scheer, U., Hügle, B., Hazan, R., and Rose, K.M. (1984). Drug-induced dispersal of transcribed rRNA genes and transcriptional products: immunolocalization and silver staining of different nucleolar components in rat cells treated with 5,6-dichloro-beta-D-ribofuranosylbenzimidazole. J. Cell Biol. 99, 672–679.

72. Yasuhara, T., Xing, Y.-H., Bauer, N.C., Lee, L., Dong, R., Yadav, T., Soberman, R.J., Rivera, M.N., and Zou, L. (2022). Condensates induced by transcription inhibition localize active chromatin to nucleoli. Mol. Cell 82, 2738–2753.e6.

73. Lindell, T.J., Weinberg, F., Morris, P.W., Roeder, R.G., and Rutter, W.J. (1970). Specific inhibition of nuclear RNA polymerase II by alpha-amanitin. Science 170, 447–449.

74. Jao, C.Y., and Salic, A. (2008). Exploring RNA transcription and turnover in vivo by using click chemistry. Proc Natl Acad Sci U S A 105, 15779–15784.

75. Riback, J.A., Eeftens, J.M., Lee, D.S.W., Quinodoz, S.A., Donlic, A., Orlovsky, N., Wiesner, L., Beckers, L., Becker, L.A., Strom, A.R., et al. (2023). Viscoelasticity and advective flow of RNA underlies nucleolar form and function. Mol Cell 83, 3095–3107.e9.

76. Bryant, C.J., McCool, M.A., Abriola, L., Surovtseva, Y.V., and Baserga, S.J. (2022). A high-throughput assay for directly monitoring nucleolar rRNA biogenesis. Open Biol 12, 210305.

77. Chao, S.H., Fujinaga, K., Marion, J.E., Taube, R., Sausville, E.A., Senderowicz, A.M., Peterlin, B.M., and Price, D.H. (2000). Flavopiridol inhibits P-TEFb and blocks HIV-1 replication. J. Biol. Chem. 275, 28345–28348.

78. Misteli, T., Cáceres, J.F., and Spector, D.L. (1997). The dynamics of a pre-mRNA splicing factor in living cells. Nature 387, 523–527.

79. Eils, R., Gerlich, D., Tvaruskó, W., Spector, D.L., and Misteli, T. (2000). Quantitative imaging of pre-mRNA splicing factors in living cells. Mol Biol Cell 11, 413–418.

80. Carvalho, T., Martins, S., Rino, J., Marinho, S., and Carmo-Fonseca, M. (2017). Pharmacological inhibition of the spliceosome subunit SF3b triggers exon junction complex-independent nonsense-mediated decay. J Cell Sci 130, 1519–1531.

81. Wu, J., Xiao, Y., Liu, Y., Wen, L., Jin, C., Liu, S., Paul, S., He, C., Regev, O., and Fei, J. (2024). Dynamics of RNA localization to nuclear speckles are connected to splicing efficiency. Sci Adv 10, eadp7727.

82. McIntyre, A.B.R., Tschan, A.B., Meyer, K., Walser, S., Rai, A.K., Fujita, K., and Pelkmans, L. (2025). Phosphorylation of a nuclear condensate regulates cohesion and mRNA retention. Nat Commun 16, 390.

83. Wei, W., Bai, L., Yan, B., Meng, W., Wang, H., Zhai, J., Si, F., and Zheng, C. (2022). When liquid-liquid phase separation meets viral infections. Front Immunol 13, 985622.

84. Li, H., Ernst, C., Kolonko-Adamska, M., Greb-Markiewicz, B., Man, J., Parissi, V., and Ng, B.W.-L. (2022). Phase separation in viral infections. Trends Microbiol 30, 1217–1231.

85. Sagan, S.M., and Weber, S.C. (2023). Let’s phase it: viruses are master architects of biomolecular condensates. Trends Biochem Sci 48, 229–243.

86. Rincheval, V., Lelek, M., Gault, E., Bouillier, C., Sitterlin, D., Blouquit-Laye, S., Galloux, M., Zimmer, C., Eleouet, J.-F., and Rameix-Welti, M.-A. (2017). Functional organization of cytoplasmic inclusion bodies in cells infected by respiratory syncytial virus. Nat Commun 8, 563.

87. Risso-Ballester, J., Galloux, M., Cao, J., Le Goffic, R., Hontonnou, F., Jobart-Malfait, A., Desquesnes, A., Sake, S.M., Haid, S., Du, M., et al. (2021). A condensate-hardening drug blocks RSV replication in vivo. Nature 595, 596–599.

88. Diot, C., Richard, C.-A., Risso-Ballester, J., Martin, D., Fix, J., Eléouët, J.-F., Sizun, C., Rameix-Welti, M.-A., and Galloux, M. (2023). Hardening of Respiratory Syncytial Virus Inclusion Bodies by Cyclopamine Proceeds through Perturbation of the Interactions of the M2-1 Protein with RNA and the P Protein. Int J Mol Sci 24. 10.3390/ijms241813862.

89. Pelletier, J., Thomas, G., and Volarević, S. (2018). Ribosome biogenesis in cancer: new players and therapeutic avenues. Nat. Rev. Cancer 18, 51–63.

90. Van Nostrand, E.L., Freese, P., Pratt, G.A., Wang, X., Wei, X., Xiao, R., Blue, S.M., Chen, J.-Y., Cody, N.A.L., Dominguez, D., et al. (2020). A large-scale binding and functional map of human RNA-binding proteins. Nature 583, 711–719.

91. Yao, R.-W., Xu, G., Wang, Y., Shan, L., Luan, P.-F., Wang, Y., Wu, M., Yang, L.-Z., Xing, Y.-H., Yang, L., et al. (2019). Nascent Pre-rRNA Sorting via Phase Separation Drives the Assembly of Dense Fibrillar Components in the Human Nucleolus. Mol Cell 76, 767– 783.e11.

92. King, M.R., Ruff, K.M., Lin, A.Z., Pant, A., Farag, M., Lalmansingh, J.M., Wu, T., Fossat, M.J., Ouyang, W., Lew, M.D., et al. (2024). Macromolecular condensation organizes nucleolar sub-phases to set up a pH gradient. Cell 187, 1889–1906.e24.

93. Kopp, K., Gasiorowski, J.Z., Chen, D., Gilmore, R., Norton, J.T., Wang, C., Leary, D.J., Chan, E.K.L., Dean, D.A., and Huang, S. (2007). Pol I transcription and pre-rRNA processing are coordinated in a transcription-dependent manner in mammalian cells. Mol Biol Cell 18, 394–403.

94. Quinodoz, S.A., Jiang, L., Abu-Alfa, A.A., Comi, T.J., Zhao, H., Yu, Q., Wiesner, L.W., Botello, J.F., Donlic, A., Soehalim, E., et al. (2025). Mapping and engineering RNA-driven architecture of the multiphase nucleolus. Nature. 10.1038/s41586-025-09207-4.

95. Donlic, A., Swanson, E.G., Chiu, L.-Y., Wicks, S.L., Juru, A.U., Cai, Z., Kassam, K., Laudeman, C., Sanaba, B.G., Sugarman, A., et al. (2022). R-BIND 2.0: An Updated Database of Bioactive RNA-Targeting Small Molecules and Associated RNA Secondary Structures. ACS Chem. Biol. 17, 1556–1566.

96. Lin, J., Zhou, D., Steitz, T.A., Polikanov, Y.S., and Gagnon, M.G. (2018). Ribosome-Targeting Antibiotics: Modes of Action, Mechanisms of Resistance, and Implications for Drug Design. Annu. Rev. Biochem. 87, 451–478.

97. Sobell, H.M. (1985). Actinomycin and DNA transcription. Proc. Natl. Acad. Sci. U. S. A. 82, 5328–5331.

98. Zhang, L., Zhang, F., Zhang, W., Chen, L., Gao, N., Men, Y., Xu, X., and Jiang, Y. (2015). Harmine suppresses homologous recombination repair and inhibits proliferation of hepatoma cells. Cancer Biol Ther 16, 1585–1592.

99. Wang, S.-K., Wu, Y., Wang, X.-Q., Kuang, G.-T., Zhang, Q., Lin, S.-L., Liu, H.-Y., Tan, J.-H., Huang, Z.-S., and Ou, T.-M. (2017). Discovery of Small Molecules for Repressing Cap-Independent Translation of Human Vascular Endothelial Growth Factor (hVEGF) as Novel Antitumor Agents. J Med Chem 60, 5306–5319.

100. Diouf, B., Lin, W., Goktug, A., Grace, C.R.R., Waddell, M.B., Bao, J., Shao, Y., Heath, R.J., Zheng, J.J., Shelat, A.A., et al. (2018). Alteration of RNA Splicing by Small-Molecule Inhibitors of the Interaction between NHP2L1 and U4. SLAS Discov 23, 164–173.

101. Pommier, Y. (2006). Topoisomerase I inhibitors: camptothecins and beyond. Nat. Rev. Cancer 6, 789–802.

102. De Isabella, P., Capranico, G., Palumbo, M., Sissi, C., Krapcho, A.P., and Zunino, F. (1993). Sequence selectivity of topoisomerase II DNA cleavage stimulated by mitoxantrone derivatives: relationships to drug DNA binding and cellular effects. Mol Pharmacol 43, 715– 721.

103. Christensen, M.O., Barthelmes, H.U., Boege, F., and Mielke, C. (2002). The N-terminal domain anchors human topoisomerase I at fibrillar centers of nucleoli and nucleolar organizer regions of mitotic chromosomes. J. Biol. Chem. 277, 35932–35938.

104. Mao, Y., Mehl, I.R., and Muller, M.T. (2002). Subnuclear distribution of topoisomerase I is linked to ongoing transcription and p53 status. Proc. Natl. Acad. Sci. U. S. A. 99, 1235– 1240.

105. Christensen, M.O., Krokowski, R.M., Barthelmes, H.U., Hock, R., Boege, F., and Mielke, C. (2004). Distinct effects of topoisomerase I and RNA polymerase I inhibitors suggest a dual mechanism of nucleolar/nucleoplasmic partitioning of topoisomerase I. J. Biol. Chem. 279, 21873–21882.

106. Staker, B.L., Hjerrild, K., Feese, M.D., Behnke, C.A., Burgin, A.B., Jr, and Stewart, L. (2002). The mechanism of topoisomerase I poisoning by a camptothecin analog. Proc Natl Acad Sci U S A 99, 15387–15392.

107. Zhang, H., Wang, J.C., and Liu, L.F. (1988). Involvement of DNA topoisomerase I in transcription of human ribosomal RNA genes. Proc Natl Acad Sci U S A 85, 1060–1064.

108. Swinney, D.C., and Anthony, J. (2011). How were new medicines discovered? Nat Rev Drug Discov 10, 507–519.

109. Donlic, A., and Hargrove, A.E. (2018). Targeting RNA in mammalian systems with small molecules. Wiley Interdiscip Rev RNA 9, e1477.

110. Tong, Y., Childs-Disney, J.L., and Disney, M.D. (2024). Targeting RNA with small molecules, from RNA structures to precision medicines: IUPHAR review: 40. Br J Pharmacol 181, 4152–4173.

111. Warner, K.D., Hajdin, C.E., and Weeks, K.M. (2018). Principles for targeting RNA with drug-like small molecules. Nat Rev Drug Discov 17, 547–558.

112. El Hage, A., French, S.L., Beyer, A.L., and Tollervey, D. (2010). Loss of Topoisomerase I leads to R-loop-mediated transcriptional blocks during ribosomal RNA synthesis. Genes Dev 24, 1546–1558.

113. Brill, S.J., DiNardo, S., Voelkel-Meiman, K., and Sternglanz, R. (1987). Need for DNA topoisomerase activity as a swivel for DNA replication for transcription of ribosomal RNA. Nature 326, 414–416.

114. Bhola, M., Abe, K., Orozco, P., Rahnamoun, H., Avila-Lopez, P., Taylor, E., Muhammad, N., Liu, B., Patel, P., Marko, J.F., et al. (2024). RNA interacts with topoisomerase I to adjust DNA topology. Mol Cell 84, 3192–3208.e11.

115. Ahmad, M., Xu, D., and Wang, W. (2017). Type IA topoisomerases can be “magicians” for both DNA and RNA in all domains of life. RNA Biol 14, 854–864.

116. Fang, L., Monroe, F., Novak, S.W., Kirk, L., Schiavon, C.R., Yu, S.B., Zhang, T., Wu, M., Kastner, K., Latif, A.A., et al. (2021). Deep learning-based point-scanning super-resolution imaging. Nat Methods 18, 406–416.

117. Beck, L.M., Shocher, A., Rossman, U., Halfon, A., Irani, M., and Oron, D. (2024). Improving correlation based super-resolution microscopy images through image fusion by self-supervised deep learning. Opt Express 32, 28195–28205.

118. Levin, B., Mittasch, M., Ferreira Gomes, B., Manteiga, J., Patel, A., Zamudio, A., Beutel, O., and Mitrea, D.M. (2021). Harnessing the power of fluorescence to characterize biomolecular condensates. In Methods in Microbiology Methods in microbiology. (Elsevier), pp. 1–47.

119. Bracha, D., Walls, M.T., and Brangwynne, C.P. (2019). Probing and engineering liquid-phase organelles. Nat Biotechnol 37, 1435–1445.

120. Ribeiro, M.T., Singh, S., and Guestrin, C. (2016). “why should I trust you?”: Explaining the predictions of any classifier. arXiv [cs.LG]. 10.48550/ARXIV.1602.04938.

121. Lee, D.S.W., Choi, C.-H., Sanders, D.W., Beckers, L., Riback, J.A., Brangwynne, C.P., and Wingreen, N.S. (2023). Size distributions of intracellular condensates reflect competition between coalescence and nucleation. Nat Phys 19, 586–596.

122. Sanders, D.W., Kedersha, N., Lee, D.S.W., Strom, A.R., Drake, V., Riback, J.A., Bracha, D., Eeftens, J.M., Iwanicki, A., Wang, A., et al. (2020). Competing Protein-RNA Interaction Networks Control Multiphase Intracellular Organization. Cell 181, 306–324.e28.

123. Stirling, D.R., Swain-Bowden, M.J., Lucas, A.M., Carpenter, A.E., Cimini, B.A., and Goodman, A. (2021). CellProfiler 4: improvements in speed, utility and usability. BMC Bioinformatics 22, 433.

124. Stringer, C., Wang, T., Michaelos, M., and Pachitariu, M. (2021). Cellpose: a generalist algorithm for cellular segmentation. Nat Methods 18, 100–106.

125. Pachitariu, M., and Stringer, C. (2022). Cellpose 2.0: how to train your own model. Nat Methods 19, 1634–1641.

126. Robinson, J.T., Thorvaldsdóttir, H., Winckler, W., Guttman, M., Lander, E.S., Getz, G., and Mesirov, J.P. (2011). Integrative genomics viewer. Nat Biotechnol 29, 24–26.

127. Quinlan, A.R., and Hall, I.M. (2010). BEDTools: a flexible suite of utilities for comparing genomic features. Bioinformatics 26, 841–842.

128. McKinley, K.L., Sekulic, N., Guo, L.Y., Tsinman, T., Black, B.E., and Cheeseman, I.M. (2015). The CENP-L-N Complex Forms a Critical Node in an Integrated Meshwork of Interactions at the Centromere-Kinetochore Interface. Mol Cell 60, 886–898.

129. Kishi, J.Y., Lapan, S.W., Beliveau, B.J., West, E.R., Zhu, A., Sasaki, H.M., Saka, S.K., Wang, Y., Cepko, C.L., and Yin, P. (2019). SABER amplifies FISH: enhanced multiplexed imaging of RNA and DNA in cells and tissues. Nat Methods 16, 533–544.

130. Ietswaart, R., Smalec, B.M., Xu, A., Choquet, K., McShane, E., Jowhar, Z.M., Guegler, C.K., Baxter-Koenigs, A.R., West, E.R., Fu, B.X.H., et al. (2024). Genome-wide quantification of RNA flow across subcellular compartments reveals determinants of the mammalian transcript life cycle. Mol Cell 84, 2765–2784.e16.

