## Supplemental Tables for "Deep Learning of Functional Perturbations from Condensate Morphology"

**Table S1. Measured $RC_{50}$ values (95% CI) in the current study and previously reported $IC_{50}$ measurements for nucleolar cap-forming compounds, related to Figure 2.** RC = response concentration, CI = confidence interval.

| | Measured $IC_{50}$ (μM) | Measurement & method | Treatment duration | Cell line | Reference |
|---|---|---|---|---|---|
| ***CX-5461*** | 0.28 - 0.38 | Nucleolar morphology response by Deep-Phase | 2 hours | U-2 OS | This work |
| | 0.15 - 0.24 | RNA intensity in DFC by 5-EU labeling | 2 hours | U-2 OS | This work |
| | 0.22 (HCT-116), 0.113 (A375), 0.054 (MIA PaCa-2) | rRNA levels by qRT-PCR | 2 hours | HCT-116, A375, MIA PaCa-2 | [1] |
| ***Act D*** | 0.0004 - 0.0014 | Nucleolar morphology response by Deep-Phase | 2 hours | U-2 OS | This work |
| | 0.002 | Metabolic labeling of rRNA intermediates resolved by gel electrophoresis | 6 hours | 2fTGH fibrosarcoma | [2] |
| ***BMH-21*** | 0.126 - 0.144 | Nucleolar morphology response by Deep-Phase | 2 hours | U-2 OS | This work |
| | 0.060 | Metabolic labeling of 47S rRNA resolved by gel electrophoresis | 4 hours | A375 | [3] |
| ***Oxaliplatin*** | 2.2 - 11.3 | Nucleolar morphology response by Deep-Phase | 2 hours | U-2 OS | This work |
| | 1.6 | Metabolic labeling of rRNA intermediates resolved by gel electrophoresis | 6 hours | 2fTGH fibrosarcoma | [2] |

**Table S2. Measured $RC_{50}$ values (95% CI) in the current study and previously reported $IC_{50}$ measurements for necklace-forming compounds, related to Figure 2.** RC = response concentration, CI = confidence interval.

| | Measured $IC_{50}$ (μM) | Measurement & method | Treatment duration | Cell line | Reference |
|---|---|---|---|---|---|
| ***FVP*** | 0.074 - 0.104 | Nucleolar morphology response by Deep-Phase | 2 hours | U-2 OS | This work |
| | 0.103 - 0.15 | RNA Pol II Phosphorylated Ser2 by antibody labeling intensity | 2 hours | U-2 OS | This work |
| | 0.13 | Metabolic labeling of rRNA intermediates resolved by gel electrophoresis | 6 hours | 2fTGH fibrosarcoma | [2] |

| | | | | | |
|---|---|---|---|---|---|
| ***THZ1*** | 0.13 - 0.18 | Nucleolar morphology response by Deep-Phase | 2 hours | U-2 OS | This work |
| | 0.1 (p-Ser2), 0.25 (p-Ser5, p-Ser7) | Western Blot against RNAPII CTD (approximated) | 4 hours | RPE-1 | [4] |
| ***DRB*** | 14 - 112 | Nucleolar morphology response by Deep-Phase | 2 hours | U-2 OS | This work |
| | 10 | Metabolic labeling of rRNA intermediates resolved by gel electrophoresis | 6 hours | 2fTGH fibrosarcoma | [2] |

**Table S3. Measured $RC_{50}$ values (95% CI) in the current study and previously reported $IC_{50}$ measurements for speckle-modulating compounds used in this study, related to Figure 3.** RC = response concentration, IC = inhibitory concentration, CI = confidence interval, N/A = not applicable.

| | Measured $IC_{50}$ (µM) | Measurement & method | Treatment duration | Cell line | Reference |
|---|---|---|---|---|---|
| ***FVP*** | 0.083 - 0.092 | Speckle morphology response by Deep-Phase | 2 hours | HEK 293T | This work |
| | 0.001 - 0.010 | Titration of purified Polymerase II and P-TEFb complex (approximated) | N/A | Pol II purified from Drosophila | [5] |
| ***Pladienolide B (Plad B)*** | 0.0023 - 0.0048 | Speckle morphology response by Deep-Phase | 2 hours | HEK 293T | This work |
| | 0.005 | Scintillation proximity assay using an immunopurified endogenous SF3B complex | 30 minutes | Complex purified from HeLa S3 nuclear extract | [6] |

**Table S4. Measured $RC_{50}$ values (95% CI) in the current study and previously reported $IC_{50}$ measurements for the RSV-modulating compound used in this study, related to Figure 3.** RC = response concentration, IC = inhibitory concentration, CI = confidence interval.

| | Measured $IC_{50}$ (µM) | Measurement & method | Treatment duration | Cell line | Reference |
|---|---|---|---|---|---|
| ***Cyclopamine (CPM)*** | 0.064 - 0.14 | RSV inclusion body (IB) morphology response by Deep-Phase | 1 hour | HEp-2 infected with RSV-M2-1-GFP-N-GFP | This work |
| | 0.064 | RSV replication inhibition in a luciferase assay | 24 hours | HEp-2 infected with RSV-Luc | [7] |

**Table S5. Fluorescence intensity values of compounds selected for screening of nucleolar morphology modulation in the 3-color nucleolar line as measured in a plate reader, related to Figure 4 and Methods.** Bolded compounds were excluded from the screen. BFP emission = 405 nm, GFP emission = 488 nm, mCherry emission = 561 nm. H = high (50 µM); M = medium (10 µM); L = low (1 µM). N/A = not applicable.

| | BFP | | | GFP | | | mCherry | | |
|---|---|---|---|---|---|---|---|---|---|
| Concentration | H | M | L | H | M | L | H | M | L |
| Ribocil-C | 108.6 | 48.6 | 127.6 | 1560.8 | 865.8 | 2184.8 | 2856.3 | 1467.3 | 3534.3 |
| Chelerythrine chloride | 125.6 | 126.6 | 124.6 | 1826.8 | 1994.8 | 2281.8 | 2958.3 | 3364.3 | 3651.3 |
| Harmine | 96.6 | 137.6 | 136.6 | 1422.8 | 2223.8 | 2245.8 | 2557.3 | 3784.3 | 3643.3 |
| **Proflavine hemisulfate** | 363.6 | 164.6 | 87.6 | 15613.8 | 4904.8 | 1628.8 | 3669.3 | 3523.3 | 2405.3 |
| Pentamidine isethionate | 124.6 | 96.6 | 129.6 | 1901.8 | 1691.8 | 2246.8 | 3194.3 | 2888.3 | 3706.3 |
| Synucleozid hydrochloride | 48.6 | 141.6 | 132.6 | 757.8 | 2219.8 | 2216.8 | 1308.3 | 3557.3 | 3631.3 |
| MIR96-IN-1 | 224.6 | 168.6 | 94.6 | 1263.8 | 1943.8 | 1613.8 | 2163.3 | 3139.3 | 2742.3 |
| SID 3712249 | 282.6 | 148.6 | 121.6 | 2283.8 | 1955.8 | 2132.8 | 3866.3 | 3420.3 | 3257.3 |
| Branaplam | 165.6 | 110.6 | 84.6 | 2458.8 | 1829.8 | 1672.8 | 4456.3 | 3224.3 | 2691.3 |
| Clomiphene citrate | 109.6 | 76.6 | 101.6 | 1701.8 | 1463.8 | 1824.8 | 3183.3 | 2466.3 | 3037.3 |
| **Furamidine dihydrochloride** | 42117.6 | 12325.6 | 1023.6 | 1638.8 | 2058.8 | 1705.8 | 3160.3 | 3528.3 | 2865.3 |
| hVEGF-IN-1 | 109.6 | 105.6 | 99.6 | 1417.8 | 1819.8 | 1701.8 | 2702.3 | 3077.3 | 2829.3 |
| Targapremir-210 | 326.6 | 164.6 | 112.6 | 2131.8 | 2250.8 | 1827.8 | 3933.3 | 3706.3 | 3145.3 |
| Myricetin | 95.6 | 107.6 | 81.6 | 1859.8 | 2174.8 | 1486.8 | 3254.3 | 3479.3 | 2414.3 |
| Risdiplam | 171.6 | 116.6 | 76.6 | 1964.8 | 1691.8 | 1331.8 | 3513.3 | 2984.3 | 2323.3 |
| Topotecan Hydrochloride | 417.6 | 1080.6 | 146.6 | 2773.8 | 4334.8 | 1562.8 | 4125.3 | 3863.3 | 2137.3 |
| MALAT1-IN-1 | 140.6 | 125.6 | 72.6 | 2089.8 | 2066.8 | 1243.8 | 3539.3 | 3459.3 | 2161.3 |
| Piperine | 199.6 | 118.6 | 66.6 | 2324.8 | 1883.8 | 1100.8 | 4164.3 | 3185.3 | 1831.3 |
| **Obatoclax Mesylate** | 91.6 | 135.6 | 62.6 | 4455.8 | 6991.8 | 2295.8 | 17397.3 | 19490.3 | 4906.3 |
| RG7800 | 140.6 | 136.6 | 85.6 | 1904.8 | 2205.8 | 1470.8 | 3306.3 | 3826.3 | 2594.3 |
| Neomycin Sulfate | 176.6 | 83.6 | 94.6 | 2477.8 | 1444.8 | 1653.8 | 4547.3 | 2385.3 | 2635.3 |
| Kanamycin Sulfate | 101.6 | 136.6 | 69.6 | 1451.8 | 2172.8 | 1246.8 | 2876.3 | 3757.3 | 2125.3 |
| Linezolid | 136.6 | 129.6 | 98.6 | 1824.8 | 2076.8 | 1601.8 | 3456.3 | 3640.3 | 2704.3 |
| DMSO | 32.6 | n/a | n/a | 756.8 | n/a | n/a | 1298.3 | n/a | n/a |
| CX-5461 | 171.6 | n/a | n/a | 2360.8 | n/a | n/a | 4095.3 | n/a | n/a |
| FVP | 159.6 | n/a | n/a | 2360.8 | n/a | n/a | 4396.3 | n/a | n/a |

**Table S6. Measured $RC_{50}$ values (95% CI) in the current study and previously reported $IC_{50}$ measurements for topoisomerase-targeting compounds used in this study, related to Figure 5.** RC = response concentration, IC = inhibitory concentration, CI = confidence interval.

| ***Drug*** | **Measured $IC_{50}$ (μM)** | **Measurement & method** | **Treatment duration** | **Cell line** | **Reference** |
|---|---|---|---|---|---|
| ***TPT*** | 1.98 - 3.10 | Nucleolar morphology response by Deep-Phase | 2 hours | U-2 OS | This work |
| | 1.43 - 3.10 | Change in anti-TOP1 nucleolar-nucleoplasmic partitioning | 2 hours | U-2 OS | This work |
| ***CPT*** | 0.9 - 2.4 | Nucleolar morphology response by Deep-Phase | 2 hours | U-2 OS | This work |

| | | | | | |
|---|---|---|---|---|---|
| | 0.8 | Metabolic labeling of rRNA intermediates resolved by gel electrophoresis | 2 hours | 2fTGH fibrosarcoma | [2] |
| ***MTX*** | 0.66 - 1.07 | Nucleolar morphology response by Deep-Phase | 2 hours | U-2 OS | This work |
| | 0.65 | Metabolic labeling of rRNA intermediates resolved by gel electrophoresis | 2 hours | 2fTGH fibrosarcoma | [2] |

**Table S7. List of screening library compounds with catalog numbers, resuspension concentrations, and resuspension medium, related to Figure 4 and Methods.**

| Compound name | Catalog # | Resuspension concentration (mM) | Resuspension medium |
|---|---|---|---|
| **Ribocil-C** | HY-19488A | 10 | DMSO |
| **Chelerythrine chloride** | HY-12048 | 10 | DMSO |
| **Harmine** | HY-N0737A | 10 | DMSO |
| **Proflavine hemisulfate** | HY-B0883 | 10 | Water |
| **Pentamidine isethionate** | HY-B0537B | 10 | DMSO |
| **Synucleozid hydrochloride** | HY-135902A | 10 | DMSO |
| **MIR96-IN-1** | HY-15843 | 10 | DMSO |
| **SID 3712249** | HY-19731 | 10 | DMSO |
| **Branaplam** | HY-19620 | 10 | DMSO |
| **Clomiphene citrate** | HY-B0463 | 10 | DMSO |
| **Furamidine dihydrochloride** | SML1559-5MG | 10 | DMSO |
| **hVEGF-IN-1** | HY-101931 | 10 | DMSO |
| **Targapremir-210** | HY-15861 | 10 | DMSO |
| **Myricetin** | HY-15097 | 10 | DMSO |
| **Risdiplam** | HY-109101 | 1 | DMSO |
| **Topotecan Hydrochloride** | HY-13768A | 10 | DMSO |
| **MALAT1-IN-1** | HY-115579 | 10 | DMSO |
| **Piperine** | HY-N0144 | 10 | DMSO |
| **Obatoclax Mesylate** | HY-10969 | 10 | DMSO |
| **RG7800** | HY-101792 | 5 | DMSO |
| **Neomycin Sulfate** | HY-B0470 | 10 | Water |
| **Kanamycin Sulfate** | HY-16566A | 10 | Water |
| **Linezolid** | HY-10394 | 10 | DMSO |

**Table S8. Oligonucleotide sequences (5'-3') used for cloning sgRNAs in HeLa TetR-Cas9 cells, related to Figure 5 and Methods.**

| | |
|---|---|
| Non-targeting sgRNA, forward sequence | CACCGAAAACAGGACGATGTGCGGC |
| Non-targeting sgRNA, reverse sequence | AAACGCCGCACATCGTCCTGTTTTC |
| TOP1-targeting sgRNA, forward sequence | CACCGTGGAAGAGGCTCATATGGTG |
| TOP1-targeting sgRNA, reverse sequence | AAACCACCATATGAGCCTCTTCCA |

**Supplemental References**


1. Drygin, D., Lin, A., Bliesath, J., Ho, C.B., O'Brien, S.E., Proffitt, C., Omori, M., Haddach, M., Schwaebe, M.K., Siddiqui-Jain, A., et al. (2011). Targeting RNA polymerase I with an oral small molecule CX-5461 inhibits ribosomal RNA synthesis and solid tumor growth. Cancer Res *71*, 1418–1430.

2. Burger, K., Mühl, B., Harasim, T., Rohrmoser, M., Malamoussi, A., Orban, M., Kellner, M., Gruber-Eber, A., Kremmer, E., Hölzel, M., et al. (2010). Chemotherapeutic drugs inhibit ribosome biogenesis at various levels. J Biol Chem *285*, 12416–12425.

3. Peltonen, K., Colis, L., Liu, H., Trivedi, R., Moubarek, M.S., Moore, H.M., Bai, B., Rudek, M.A., Bieberich, C.J., and Laiho, M. (2014). A targeting modality for destruction of RNA polymerase I that possesses anticancer activity. Cancer Cell *25*, 77–90.

4. Kwiatkowski, N., Zhang, T., Rahl, P.B., Abraham, B.J., Reddy, J., Ficarro, S.B., Dastur, A., Amzallag, A., Ramaswamy, S., Tesar, B., et al. (2014). Targeting transcription regulation in cancer with a covalent CDK7 inhibitor. Nature *511*, 616–620.

5. Chao, S.H., and Price, D.H. (2001). Flavopiridol inactivates P-TEFb and blocks most RNA polymerase II transcription in vivo. J Biol Chem *276*, 31793–31799.

6. Cretu, C., Agrawal, A.A., Cook, A., Will, C.L., Fekkes, P., Smith, P.G., Lührmann, R., Larsen, N., Buonamici, S., and Pena, V. (2018). Structural Basis of Splicing Modulation by Antitumor Macrolide Compounds. Mol Cell *70*, 265–273.e8.

7. Risso-Ballester, J., Galloux, M., Cao, J., Le Goffic, R., Hontonnou, F., Jobart-Malfait, A., Desquesnes, A., Sake, S.M., Haid, S., Du, M., et al. (2021). A condensate-hardening drug blocks RSV replication in vivo. Nature *595*, 596–599.
